# Negative Frequency-dependent Mimicry Governs Seasonal Population Dynamics of Batesian Mimics

**DOI:** 10.64898/2026.08.05.743083

**Authors:** Viraj Nawge, Rohit Girotra, Nagesh Ramamurthy, Krushnamegh Kunte

## Abstract

Seasonal changes in climate, resources, and trophic interactions jointly shape prey population dynamics in ecological systems. In monsoon-driven tropical and subtropical landscapes, prey populations cycle between cool, wet, favourable periods during the rainy seasons and hot/cold, dry and sub-optimal conditions outside the rainy seasons, with resource availability and predation risk changing across seasons of the year. Defensive strategies and species interactions such as aposematism and Batesian mimicry are expected to interact with seasonal changes in resource availability and predation risk in determining population dynamics of prey species. Here we study population dynamics of mimetic butterfly community members using a 12-year long-term dataset from a subtropical urban forest in peninsular India. Our results show that climate and species interactions differentially influence population dynamics of different functional categories in mimetic butterfly communities, i.e., of aposematic species, mimetic and non-mimetic forms of mimetic species, and close relatives treated as ecological and phylogenetic contrasts. Population dynamics of non-mimetic species and forms were predominantly influenced by climate parameters such as temperature and precipitation, whereas population dynamics of mimics were more deeply impacted by mimetic interactions. Population dynamics of non-mimetic and mimetic forms of the same species showed distinct decoupling, with population dynamics of non-mimetic forms being similar to their non-mimetic relatives (phylogenetic contrasts). On the other hand, population dynamics of aposematic species and mimetic forms/species followed the predictions of negative frequency dependence and phase-shifting in mimicry theory: (a) mimetic forms/species were less abundant than their Batesian models, (b) the harmonic mean of populations of Batesian models influenced the upper limit of relative frequency of mimetic forms/species to a greater degree in these continuously breeding, seasonally fluctuating populations, and (c) populations of Batesian mimics peaked after population peaks of their Batesian models. These results reveal that climate and species interactions differentially determine population dynamics of prey species by functional categories at the community level, rather than by species identity and individual species attributes and resource demands.

## INTRODUCTION

Many tropical and subtropical habitats lack the harsh thermal extremes that induce dormancy and/or long periods of relative inactivity in temperate regions. Instead, most species remain active year-round, albeit at varying population sizes (Bonebrake et al. 2010). In these areas many invertebrates breed continuously, resulting in overlapping generations and complex seasonal dynamics driven largely by climate and resource availability. Rather than a clear dichotomy between “active” and “dormant” seasons, tropical systems are better characterized by a suitability spectrum ranging from optimal to sub-optimal conditions (Kishimoto Yamada and Itioka 2015; Wolda 1988). In peninsular India, this spectrum aligns strongly with the wet–dry monsoon cycle: the cold, dry winters and hot, dry summers are sub-optimal for insect activity, whereas rainy seasons bring cooler, wetter conditions and higher humidity that create more favourable climatic and resource regimes for adult flight and immature survival (hypothesis “H_1_: Climate”, in Fig. 1A).

**Figure 1.**
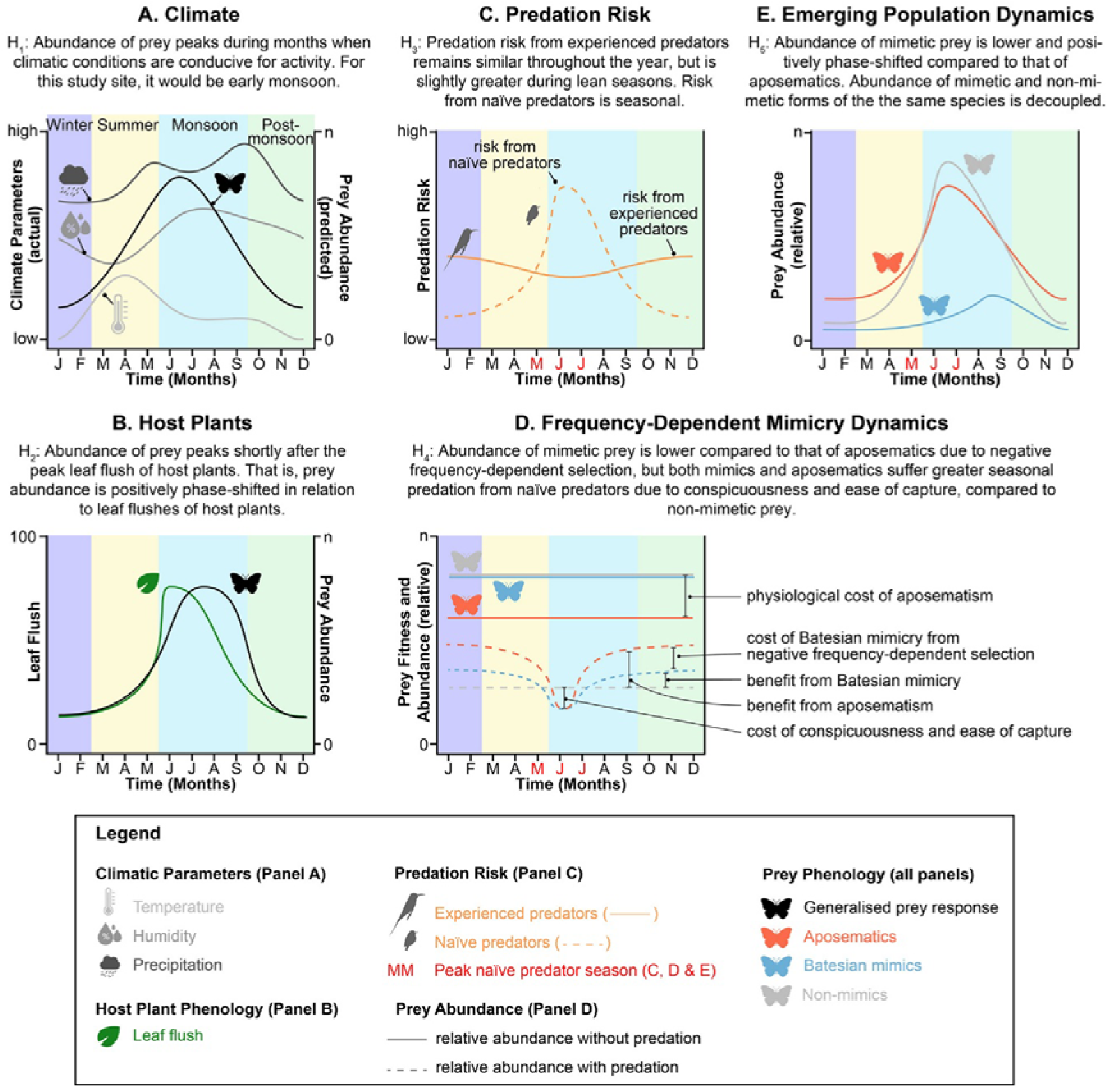
Conceptual framework illustrating multiple ecological selection pressures that shape prey population dynamics. **(A) Climate**: Seasonally fluctuating climatic factors constrain prey abundance to a limited optimal period for activity. **(B) Host plants**: Seasonal availability of larval host plants determines the timing of prey activity, leading to either close overlap or a positive phase shift between resource and prey peaks. **(C) Predation risk**: Experienced and naïve predators impose different predation risks across the year, with naïve predators concentrated in the breeding season. **(D) Frequency-dependent mimicry dynamics**: Naïve predators impose disproportionate predation risk on aposematic species and their Batesian mimics; once predators are experienced, these defended forms gain an advantage over non-mimetic prey, but mimic fitness is constrained by negative frequency-dependent selection. **(E) Emerging population dynamics**: Under differential predation risk, mimetic prey maintains lower abundances than models and may be temporally phase-shifted to avoid periods of high naïve predation. In sex-limited mimicry systems, only mimetic forms are strongly constrained by frequency-dependent selection, whereas non-mimetic forms may be governed more by climate or host-plant dynamics, resulting in decoupled population trajectories within a species.

Within this seasonal climate, the phenology of food plant resources imposes a key bottom-up constraint on prey life cycles. In many plant-consumer systems, trees and shrubs are evergreen or semi-evergreen but shift resource allocation between foliage and reproduction across the year, creating seasonal pulses of relatively high-quality food (Datta et al. 2025; Navarro-Cano et al. 2015; Peterson 1997; Toftegaard et al. 2019). Annual plants, in contrast, show tight phenological coupling with rainfall, often germinating and flowering during a narrow period of the rainy season. For herbivorous prey, this generates a predictable temporal sequence from leaf flush to larval development to adult emergence, resulting in lagged peaks in adult abundance (H_2_: Host plant, Fig. 1B). Comparable patterns across ecosystems show close matching between resource phenology, predation pressure, and prey abundance, underscoring the importance of resource-driven lags in shaping population trajectories (Both et al. 2009; Getman-Pickering et al. 2023; Murali and Sukumar 1993; Posledovich et al. 2018; Toftegaard et al. 2019).

At the same time, the abundance and experience of predators may impose top-down selection on population sizes and phase-shifting in prey population dynamics. In many (sub)tropical systems, predators breed during the hot, dry season, when food resources are relatively scarce. Adults may increase foraging effort to support reproduction, and later to provision growing offspring, such that the period of hungry, naïve juveniles coincides with increasing prey abundance after the onset of the rainy season (Stouffer, Johnson, and Bierregaard 2013). Naïve predators are more likely to attack conspicuous or novel prey that are easy to detect and attack (Mappes et al. 2014; Zvereva and Kozlov 2021), which can impose strong seasonal selection on aposematic species whose warning signals depend on predator learning (Fig.1C). While edible prey experience relatively constant predation pressure, aposematic species face episodic peaks of naïve predation that can both exact short-term costs and provide long-term benefits through initial predator learning (Fig. 1D). Batesian mimics, which are palatable to predators but resemble defended models as a defence strategy, benefit only when they remain rare relative to Batesian models and/or emerge after the naïve predators have been educated. Therefore, their fitness is expected to be tightly coupled with model abundance and predator experience (Fig.1D) (Brisson 2018; Kunte 2009; Kunte, Kizhakke, and Nawge 2021). Under negative frequency-dependent selection, mimetic forms should be less abundant (Ries and Mullen 2008) and may also be temporally offset from defended models to avoid exposing naïve predators to frequent mimic encounters (Kunte et al. 2021; Waldbauer 1988).

These interacting forces – climate-driven suitability gradients, lagged responses to resource phenology, and seasonally fluctuating predation risk in a frequency-dependent mimicry landscape – are expected to generate distinct emergent population dynamics for prey (Fig.1E). Mimetic prey should be less abundant and positively phase-shifted with respect to defended models, and, in sexually dimorphic or polymorphic species, the phenology of mimetic and non-mimetic forms may become partially decoupled. For example, in systems where only some individuals or one sex adopts a mimetic strategy, mimetic individuals may have a higher fitness due to their relationships but may be constrained to a narrower seasonal window and/or resources imposed by the negative frequence dependence in Batesian mimicry, while non-mimetic forms exploit broader resource and risk landscapes. In such cases, the rarity of mimetic forms emerges not from static “optimal” frequencies but from ongoing feedbacks among predators, Batesian models, and mimics.

Based on this background, we formulated the following hypotheses (Kunte et al. 2021), which are illustrated in Fig. 1:

H_1_, based on seasonal climatic fluctuations (abiotic factors): Abundance of prey peaks during months when climatic conditions are conducive for activity of cold-blooded insects. For our study site, it would be early monsoon.

H_2_, based on seasonal availability of larval host plants (bottom-up effects): Abundance of prey peaks shortly after the peak leaf flush of host plants. That is, prey abundance is positively phase-shifted in relation to leaf flushes of host plants.

H_3_, based on seasonal variation in predation risk (top-down effects): Predation risk from experienced predators remains similar throughout the year, but is slightly greater during lean seasons. Risk from naïve predators is seasonal.

H_4_, based on negative frequency-dependent selection on Batesian mimicry (defensive strategies and species interactions): Abundance of mimetic prey is lower compared to that of their Batesian models (aposematic species) due to negative frequency-dependent selection, but both mimics and aposematics suffer greater seasonal predation from naïve predators due to conspicuousness and ease of capture, compared to non-mimetic prey.

H_5_, based on emerging population dynamics in mimicry systems (H_1_-H_4_ combined): Abundance of mimetic prey is lower and positively phase-shifted compared to that of the aposematic species. Abundance of mimetic and non-mimetic forms of the the same species is decoupled. This hypothesis may be further subdivided for clarity as follows: H_5a_: Abundance of mimetic prey is lower compared to that of aposematic species due to negative frequency-dependent selection on Batesian mimicry. H_5b_: Abundance of mimetic prey is positively phase-shifted compared to that of aposematic prey, after naïve predators have been educated. H_5c_: Seasonal abundance of mimetic and non-mimetic forms of the the same species is decoupled because of mimicry.

Here we test these hypotheses using a 12 year time series of butterfly population data from a subtropical urban forest in peninsular India. The study site supports a rich and diverse butterfly community, including multiple species with aposematic and Batesian mimetic strategies. Several of these species occur in di/polymorphic forms, providing a natural setting to examine how climate, resource phenology, and frequency dependent mimicry jointly shape abundance peaks, temporal phase shifts, and intraspecific population decoupling in continuously breeding populations. By focusing on butterfly phenology and mimicry, we illustrate how general principles of prey–resource–predator interactions give rise to distinct population dynamics in seasonally structured environments.

## MATERIALS AND METHODS

**a. Study area and butterfly surveys:** We conducted the study at the Doresanipalya Forest Research Station (JP Nagar Reserve Forest) in Bengaluru, Karnataka, India, a protected urban forest. We surveyed butterflies fortnightly since 2012 as part of the Indian Butterfly Monitoring Scheme (iBMS, https://www.ibms-network.in/), with a six-month interruption during the COVID-19 restrictions, yielding a 12-year time series of counts. On each survey day we carried out six consecutive 30-min time-constrained counts between 09.00 and 12.00 hours along existing trails, recording all butterflies detected within a practical observation distance (Attiwilli, Ravikanthachari, and Kunte 2024). Experienced volunteers from Bengaluru Butterfly Club (BBC) helped with the surveys. Species were identified using field guides, collective expertise, and photographic records, and uncertain identifications were resolved by group consensus or later verification. We compiled citizen science data for the same species from the Butterflies of India website (https://www.ifoundbutterflies.org; (Kunte, Sondhi, and Roy 2026) and Global Biodiversity Informatics Facility (GBIF; https://www.gbif.org).
**b. Study system: mimetic communities and their phylogenetic and ecological contrasts:** We focused on the well-defined mimetic butterfly communities from peninsular India (Basu, Bhaumik, and Kunte 2023; Joshi, Prakash, and Kunte 2017; Su, Lim, and Kunte 2015). The study site supports six of the known seven mimetic communities (Fig. S1), but with fewer species per ring than in wet evergreen forests of the Western Ghats that includes a few endemics. The mimetic communities comprised several aposematic models and their Batesian mimics. We identified phylogenetic contrasts of mimetic species from closely related non-mimetic taxa (congeners or members of sister clades) to distinguish mimicry- specific effects from shared ancestry (Basu et al. 2023; Joshi et al. 2017). Ecological contrasts of aposematic species included single aposematic species that did not have any Batesian mimics. These contrasts tested whether climate and frequency/density-dependent mimicry uniquely shaped mimetic rarity and population dynamics.
**c. Climate data:** We characterized local climatic conditions using reanalysis-based temperature and humidity data along with gridded precipitation data for the grid cell (0.25°) encompassing the study site. Air temperature and dew point temperature were extracted from the ERA5 hourly reanalysis dataset (Hersbach et al. 2023) for the full study period, and relative humidity was derived from these variables. Daily temperature and humidity metrics were aggregated to monthly scales to match butterfly count data (Fig. S2). Precipitation was obtained from the India Meteorological Department (IMD) daily gridded rainfall dataset (Pai et al. 2014) and aggregated to monthly totals for analysis.
**d. Host plant data and phenology:** Larval host plants for each focal and contrast butterfly species were compiled from the continually updated Butterflies of India website. For each butterfly species, we extracted documented larval hosts and restricted this list to plant species occurring in peninsular India using the Digital Flora of Peninsular India database (https://indiaflora-ces.iisc.ac.in/FloraPeninsular/faq.php). Flowering and fruiting phenology for these host plants was obtained from the same database, based on herbarium records and field observations reporting typical months of reproductive activity. These phenology data served as a qualitative indicator of peak resource availability rather than a detailed demographic model of host plants (Fig.S4). Leafing season was assumed to be two or three months prior to flowering and fruiting as per the seasonal climatic cycles and plant phenology.
**e. Bird breeding phenology:** List of insectivorous birds occurring at the study site was compiled from eBird and filtered for bird species that are known and potential predators of butterfly prey species (Sullivan et al. 2009). Breeding phenology of birds was extracted from ornithological books (Ali and Ripley 1983), and occurrence records for the greater Bengaluru area were obtained from the citizen-science dataset available on GBIF (Fig. S5).
**f. Data processing and statistical analyses:** All data compilation and analyses were conducted in R v4.3.3 (v.4.3.2; R Core Team, R Foundation for Statistical Computing, Vienna, Austria). We used generalized additive models (Wood 2011), *rpart* for recursive partitioning and regression trees (Breiman et al. 1984), *mgcv* for generalized additive models, and circular for circular statistics (Fisher 1993). Data handling and visualization were aided by standard *tidyverse* tools where appropriate.
**i. Standardization of counts and relative abundance:** Raw 30-min survey counts were aggregated to monthly totals and standardized to a common sampling effort of 10 count slots per month. Months with fewer than six completed slots were excluded to avoid unreliable estimates based on very low effort. Standardized monthly counts were analysed using generalized additive models(Edwards and Crone 2021; Wood 2011) with a negative binomial error distribution and log link, with species, month, and year included to account for differences in abundance while controlling for temporal variation.
**ii. Phenology and activity peaks:** Species activity peaks were estimated from fitted GAM-based flight curves. The month of maximum fitted abundance was taken as the peak activity month, and uncertainty was assessed by bootstrapping. Overlap in peak activity among species was also estimated from the bootstrap distributions. Circular statistics were used to compare peak timing among species.
**iii. Climate–count relationships:** To examine how short-term climatic variation influenced butterfly counts, we used both tree-based recursive partitioning and generalized additive models. For the tree-based analysis, we fitted regression trees using *rpart* with a Poisson loss function. We considered five climatic predictors: monthly minimum temperature, maximum temperature, mean temperature, total precipitation, and mean relative humidity. Climate variables were aggregated to monthly values to retain fine-scale temporal variation and avoid artifacts from coarser temporal averaging. The tree models were fitted separately for the five functional categories.

We pooled counts across functional categories and regressed total monthly counts against continuous monthly climate variables. This pooling of data did not change the variance explained estimation much, evident form difference between variance explained by first split for individual species and the group (Fig. S9). The resulting full trees were pruned using the based on the cross-validation error, where a tree with highest complexity parameter within one standard deviation on minimum cross validation error tree was selected, and we extracted the primary split points (“critical values”) for each climatic variable from the final trees. Variance explained for each tree were derived from node deviances. 95% confidence on the variance explained was derived by bootstrapping and used a bootstrap test to compare differences.

To corroborate and generalize the climate–count relationships identified by the tree models, we additionally fitted generalized additive models to monthly counts of functional categories using *mgcv*. In these models, monthly counts were regressed on the same set of climatic covariates: count_i_∼f_1_(Tmin_i_)+f_2_(Tmax_i_)+f_3_(Tmean_i_)+f_4_(precipitation_i_)+f_5_(RH_i_) f_1_…f_5_ are smooth functions estimated using penalized regression splines. Model terms with weak support were removed sequentially. Concurvity was assessed using the *mgcv*, *concurvity* function, and interaction terms were considered for variables with high concurvity. Variables showing approximately linear relationships were confirmed using effective degrees of freedom from *gam.check*, and linear effects were included as parametric terms where appropriate.

This combination of recursive partitioning and GAMs allowed us to identify both sharp climatic thresholds and more gradual, potentially nonlinear relationships between climate and butterfly abundance.

## RESULTS

### a. Climatic factors explained significant seasonal variation in population abundance of non-mimetic and aposematic butterflies, but not of the Batesian mimics

Recursive partitioning trees identified clear climatic thresholds, statistically supported by minimum cross-validation error, that governed the effects of each climatic variable on the abundance of different functional categories within the mimetic communities and phylogenetic/ecological contrasts. Overall, climatic factors significantly explained seasonal variation in population abundance of Batesian models (i.e., aposematic species) and non-mimetic butterflies (i.e., non-mimetic forms of mimetic species, and non-mimetic sister species/groups that we treated as phylogenetic contrasts of Batesian mimics) (variation explained: aposematics: 24.4% (95% CI: 18.38 – 39.79); non-mimetic forms of mimetic species: 31.37% (95% CI: 21.02 – 33.08); and phylogenetic contrasts: 18.34% (95% CI: 12.09 – 31.67) (Fig. 2, Table 2). This pattern was even more pronounced at the level of individual species (Fig. S3).

**Figure 2.**
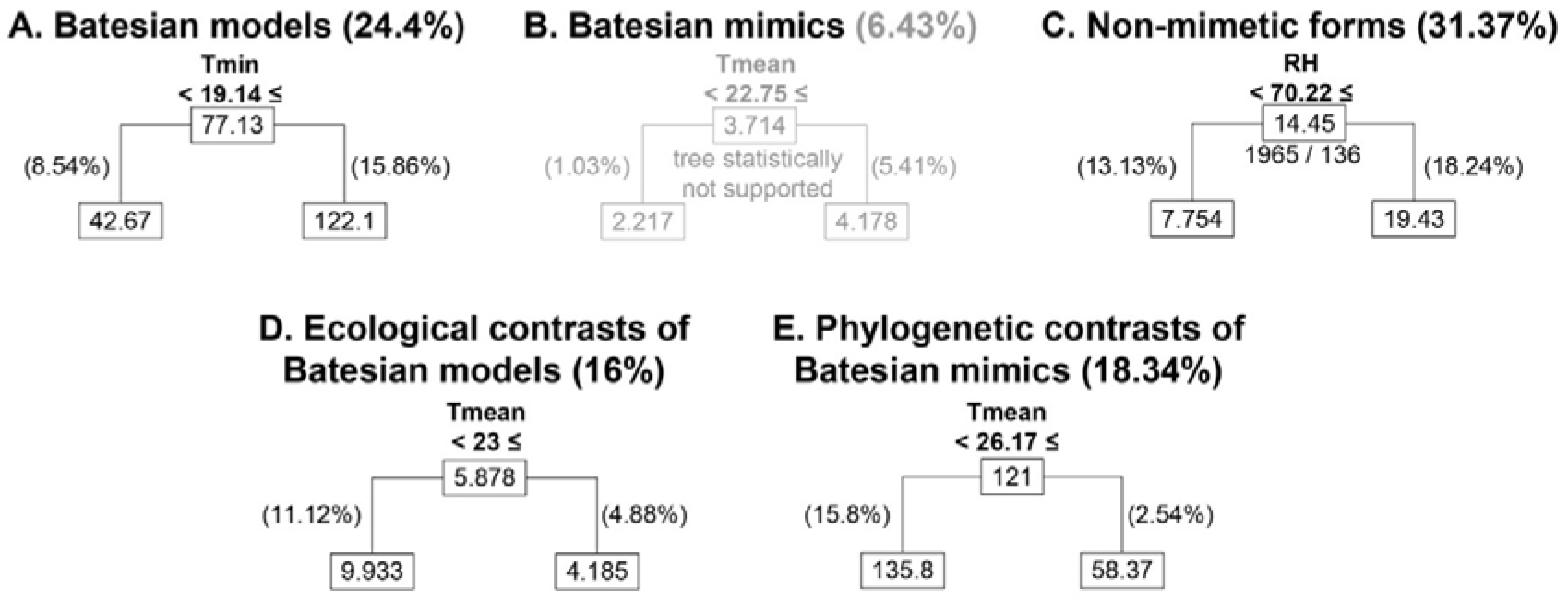
Climatic factors explain variance in seasonal abundance of aposematic and non-mimetic butterflies, but not of Batesian mimics. Recursive partitioning tree models, pruned at the minimum cross validation error for each functional category, show that counts of Batesian mimics lack statistically supported splits with respect to any climatic variable. For other groups, the value at the root node indicates the critical threshold for the first split along the respective climatic parameter; the values in the boxes are average normalized monthly counts for that node. Tmin: minimum temperature, Tmean: mean temperature, RH: relative humidity. The values in brackets next to each functional category represent the total percentage of variance explained by the tree, and the values on the branches indicate the percentage of variance explained by that branch. See Fig. S7 for the maximized, unpruned trees.

**Table 1.** Effect of climatic variable on each functional category. Positive (+ve) impact indicates proportional effect and negative impact (-ve) indicates inversely proportion effect.

| Climate parameter | Functional category | Threshold value | Impact on abundance | Variance explained (%) |
| --- | --- | --- | --- | --- |
| Minimum temperature (°C) | Batesian models | 19.14 | +ve | 24.41 |
|  | Batesian mimics | 15.81 | +ve | 5.98 |
|  | Non-mimetic forms | 17.14 | +ve | 8.74 |
|  | Ecological contrasts of Batesian models | 13.12 | -ve | 6.73 |
|  | Phylogenetic contrasts of Batesian mimics | 20.53 | -ve | 13.1 |
| Mean temperature (°C) | Batesian models | 23.84 | +ve | 15.24 |
|  | Batesian mimics | 22.75 | +ve | 6.45 |
|  | Non-mimetic forms | 26.43 | -ve | 15.28 |
|  | Ecological contrasts of Batesian models | 23 | -ve | 16 |
|  | Phylogenetic contrasts of Batesian mimics | 26.17 | -ve | 18.34 |
| Maximum temperature (°C) | Batesian models | 31.24 | +ve | 6.12 |
|  | Batesian mimics | 34.9 | -ve | 4.35 |
|  | Non-mimetic forms | 32.7 | -ve | 14.46 |
|  | Ecological contrasts of Batesian models | 29.14 | -ve | 11.93 |
|  | Phylogenetic contrasts of Batesian mimics | 33.5 | -ve | 14.87 |
| Relative humidity (%) | Batesian models | 68.85 | +ve | 6.58 |
|  | Batesian mimics | 56.54 | +ve | 3.71 |
|  | Non-mimetic forms | 70.22 | +ve | 31.37 |
|  | Ecological contrasts of Batesian models | 60.67 | +ve | 5.6 |
|  | Phylogenetic contrasts of Batesian mimics | 57.01 | +ve | 8.96 |
| Precipitation (mm) | Batesian models | 13.76 | +ve | 18.8 |
|  | Batesian mimics | 8.71 | +ve | 5.83 |
|  | Non-mimetic forms | 3.63 | +ve | 17.11 |
|  | Ecological contrasts of Batesian models | 2.35 | +ve | 4.24 |
|  | Phylogenetic contrasts of Batesian mimics | 24.08 | -ve | 4.83 |

**Table 2.** Variance in butterfly abundance explained by climatic factors. Bold values indicate trees supported with minimum cross-validation error. Confidence intervals were calculated by bootstrapping. **B.** Statistical comparison for variance explained by climate tree models. It compares trees generated by bootstrapping across functional categories (Kruskal- Wallis χ^2^(df) = 2683.9(7), p-value < 0.0001). Pairwise results are shown for Dunn’s tests with Bonferroni correction for multiple comparisons.

**A.**
| **Functional category** | **Variance explained (%)** | **SE** | **95% CI** |
| --- | --- | --- | --- |
| Batesian models | **24.4** | 5.46 | 18.38 – 39.79 |
| Batesian mimics | 6.43 | 3.04 | 6.24–18.14 |
| Non-mimetic forms | **31.37** | 6.07 | 21.02–33.08 |
| Ecological contrasts of Batesian models | **16** | 6.46 | 7.76–33.08 |
| Phylogenetic contrasts of Batesian mimics | **18.34** | 4.99 | 12.09–31.67 |
| Abundance control 1 | **32.41** | 6.71 | 13.18–39.48 |
| Abundance control 2 | **21.89** | 4.26 | 16.3–33.01 |
| Abundance control 3 | **36.04** | 4.93 | 32.21–51.52 |

**B.**
| **Functional category** | **Pairwise test** |  |
| --- | --- | --- |
|  | **Z** | **p** |
| Batesian mimics vs Non-mimetic forms | -34.6714 | < **0.0001** |
| Batesian mimics vs Phylogenetic contrasts | -16.2901 | < **0.0001** |
| Batesian models vs Ecological contrasts | 15.2133 | < **0.0001** |
| Batesian models vs Batesian mimics | 27.8801 | < **0.0001** |
| Batesian models vs Non-mimetic forms | -6.79129 | < **0.0001** |
| Non-mimetic forms vs Phylogenetic contrasts | 18.3813 | < **0.0001** |
| Batesian mimics vs Abundance control 1 | -23.913 | < **0.0001** |
| Batesian mimics vs Abundance control 2 | -21.153 | < **0.0001** |
| Batesian mimics vs Abundance control 3 | -45.304 | < **0.0001** |

Relative humidity and precipitation showed broadly positive effects, with higher values of these parameters associated with increased butterfly abundance (Table 1). Phylogenetic contrasts indicated a negative effect of precipitation, although the variance explained was small. Precipitation appeared weakly informative, as the identified thresholds were low relative to the full range of observed precipitation values (Table 1, Fig. S2). Minimum and mean temperatures exerted a strong influence on population abundance, although the exact effects varied across functional categories, and maximum temperature had a negative effect on abundance (Table 1).

In contrast, climatic parameters poorly explained seasonal variation in population abundance of Batesian mimics. (variation explained: 6.43% (95% CI: 6.24–18.14), with no statistical support from the analysis of cross-validation error; Table 2, Fig. 2). Since Batesian mimics were low in abundance compared to aposematic and non-mimetic butterflies, we tested the effects of low abundance on the statistical inference, by comparing an unrelated set of non-mimetic species that had seasonal abundance similar to that of the Batesian mimics. This comparison revealed that the lack of statistical support for the low variance explained in Batesian mimics was not due to their low abundance: climatic parameters explained a high proportion of seasonal variation in low-abundance non-mimetic species (i.e., abundance controls) (variation explained: 21–36%; Table 2). Differences in seasonal population abundance with respect to climatic variables among all the functional groups discussed above were statistically well supported (Table 2). Notably, primary climatic thresholds differed markedly between mimetic and non-mimetic butterflies (Fig. S3), with thresholds for non-mimetic butterflies aligning closely with those of their phylogenetic contrasts, particularly for temperature.

These patterns were further corroborated by generalized additive model (GAM) analyses, which detected statistically significant but relatively weak climatic effects for Batesian mimics, explaining 17% of the deviance (Table 3, Fig. 3). By their statistical nature, the tree-based models are inherently parsimonious and capture strong, threshold-like responses, whereas GAMs are better suited to detect smooth and gradual relationships. Thus, with both these types of analysis, the above results collectively suggest that climatic variables played strong, statistically supported roles in regulating seasonal population abundance of aposematic and non-mimetic butterflies, but not of the Batesian mimics.

**Figure 3.**
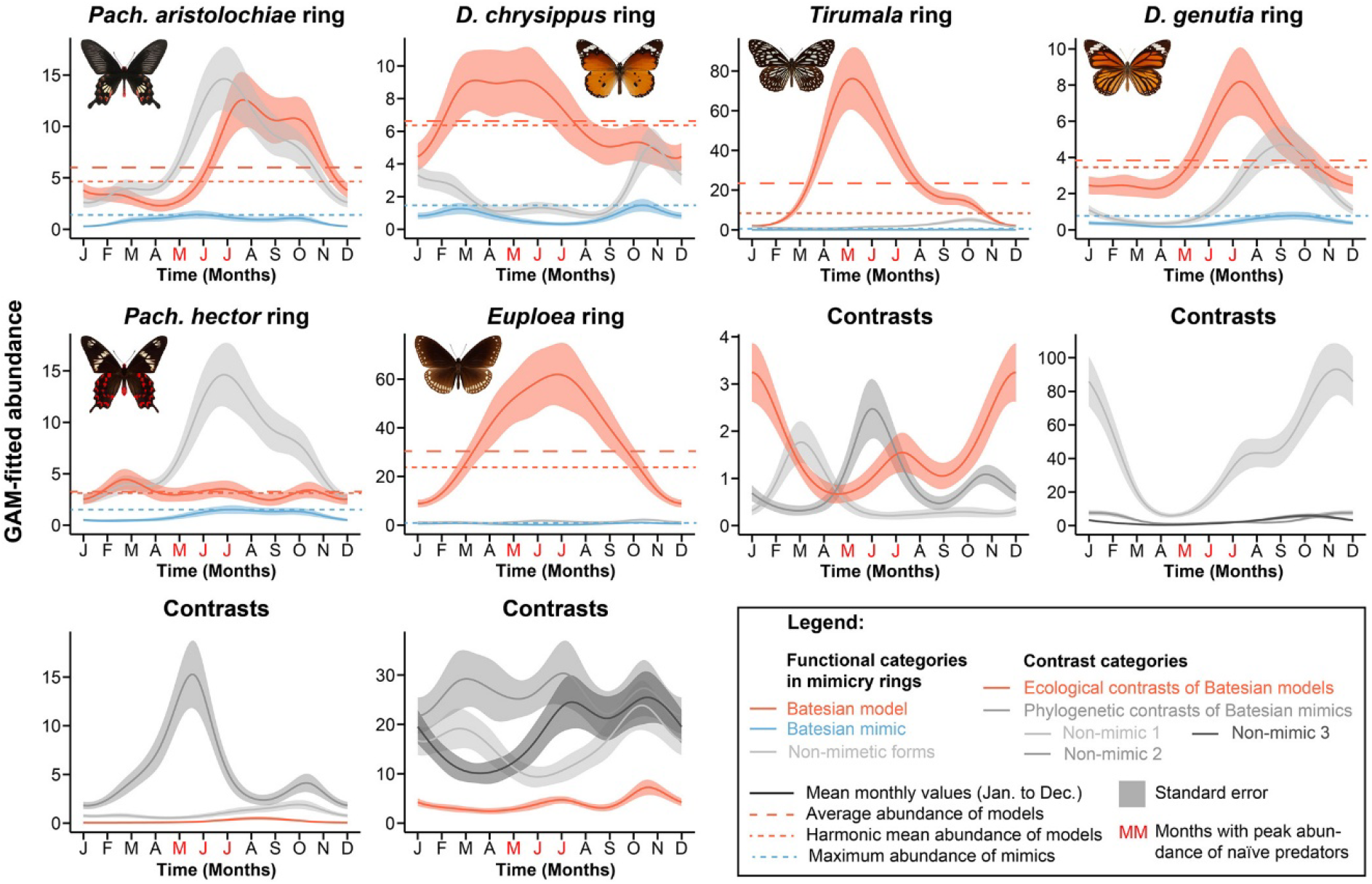
Harmonic mean determines the frequency-dependent limit for mimics in continuously breeding prey populations. Generalized additive model (GAM)–fitted abundance curves for species in various mimicry rings and their phylogenetic and ecological contrasts are shown. In mimicry-ring plots, the dashed line represents the average abundance of the Batesian model, while the dotted line indicates its harmonic mean abundance. For Batesian mimics, the dotted line represents maximum observed abundance. For a complete species legend, see Fig. S6.

**Table 3A.**
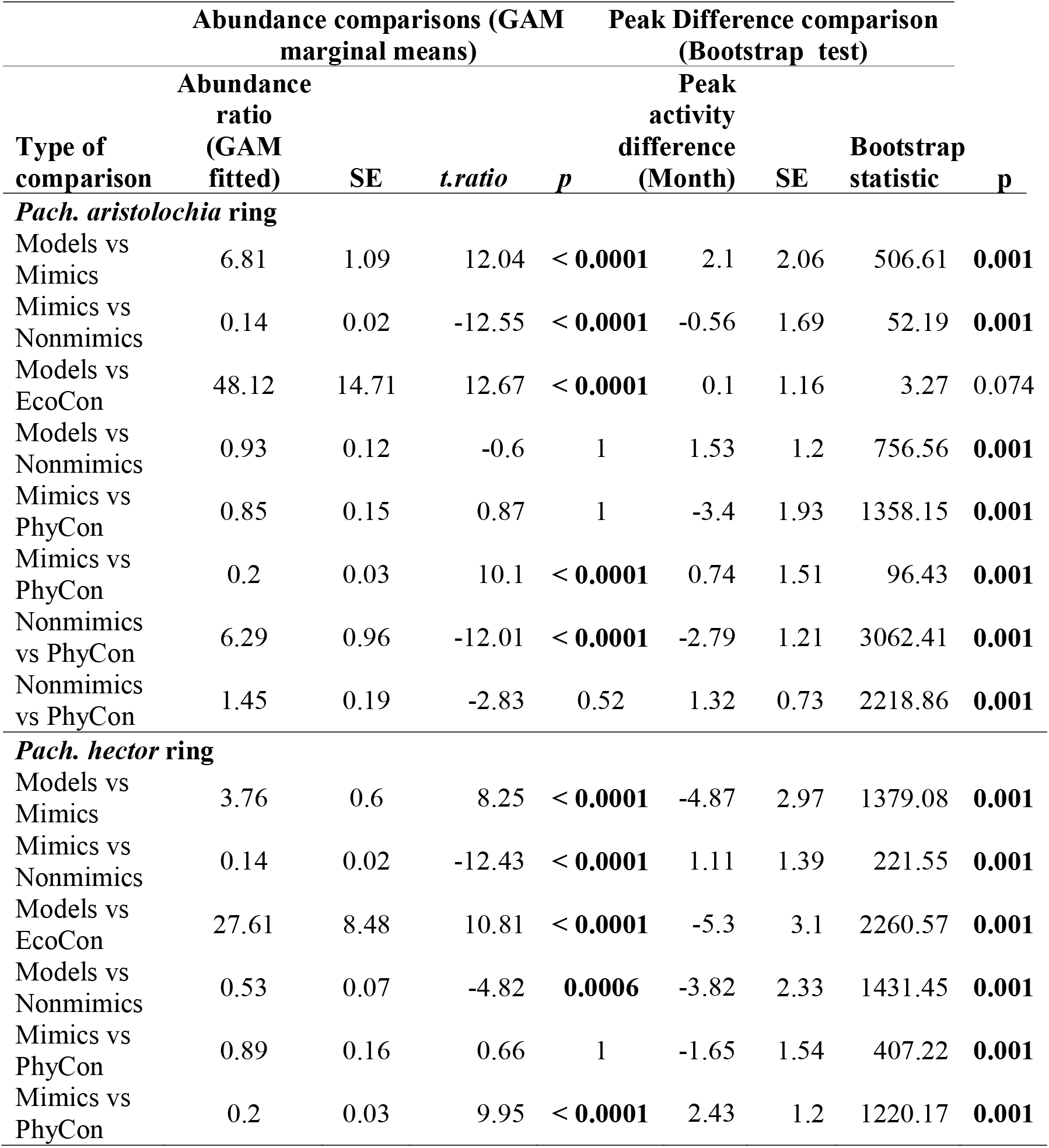
Comparison of seasonal population abundance across functional categories of mimetic butterfly communities and their phylogenetic and ecological contrasts. Abundance ratio obtained through marginal means of fitted GAM models. Peak activity difference is obtained by mean of pairwise differences of bootstrapped replicate. Functional groups are shortened for better readability: Models: Batesian models, Mimics: Batesian mimics, Nonmimics: Non-mimetic forms of Batesian mimics, EcoCon: Ecological contrasts of Batesian models, PhyCon: Phylogenetics contrasts of Batesian mimics.

| Type of comparison | Abundance comparisons (GAM marginal means) |  |  |  | Peak Difference comparison (Bootstrap test) |  |  |  |
| --- | --- | --- | --- | --- | --- | --- | --- | --- |
|  | Abundance ratio (GAM fitted) | SE | *t*-ratio | *p* | Peak activity difference (Month) | SE | Bootstrap statistic | *p* |
| ***Pach. aristolochia* ring** |  |  |  |  |  |  |  |  |
| Models vs Mimics | 6.81 | 1.09 | 12.04 | < **0.0001** | 2.1 | 2.06 | 506.61 | **0.001** |
| Mimics vs Nonmimics | 0.14 | 0.02 | -12.55 | < **0.0001** | -0.56 | 1.69 | 52.19 | **0.001** |
| Models vs EcoCon | 48.12 | 14.71 | 12.67 | < **0.0001** | 0.1 | 1.16 | 3.27 | 0.074 |
| Models vs Nonmimics | 0.93 | 0.12 | -0.6 | 1 | 1.53 | 1.2 | 756.56 | **0.001** |
| Mimics vs PhyCon | 0.85 | 0.15 | 0.87 | 1 | -3.4 | 1.93 | 1358.15 | **0.001** |
| Mimics vs PhyCon | 0.2 | 0.03 | 10.1 | < **0.0001** | 0.74 | 1.51 | 96.43 | **0.001** |
| Nonmimics vs PhyCon | 6.29 | 0.96 | -12.01 | < **0.0001** | -2.79 | 1.21 | 3062.41 | **0.001** |
| Nonmimics vs PhyCon | 1.45 | 0.19 | -2.83 | 0.52 | 1.32 | 0.73 | 2218.86 | **0.001** |
| ***Pach. hector* ring** |  |  |  |  |  |  |  |  |
| Models vs Mimics | 3.76 | 0.6 | 8.25 | < **0.0001** | -4.87 | 2.97 | 1379.08 | **0.001** |
| Mimics vs Nonmimics | 0.14 | 0.02 | -12.43 | < **0.0001** | 1.11 | 1.39 | 221.55 | **0.001** |
| Models vs EcoCon | 27.61 | 8.48 | 10.81 | < **0.0001** | -5.3 | 3.1 | 2260.57 | **0.001** |
| Models vs Nonmimics | 0.53 | 0.07 | -4.82 | **0.0006** | -3.82 | 2.33 | 1431.45 | **0.001** |
| Mimics vs PhyCon | 0.89 | 0.16 | 0.66 | 1 | -1.65 | 1.54 | 407.22 | **0.001** |
| Mimics vs PhyCon | 0.2 | 0.03 | 9.95 | < **0.0001** | 2.43 | 1.2 | 1220.17 | **0.001** |

**Table 3B.**
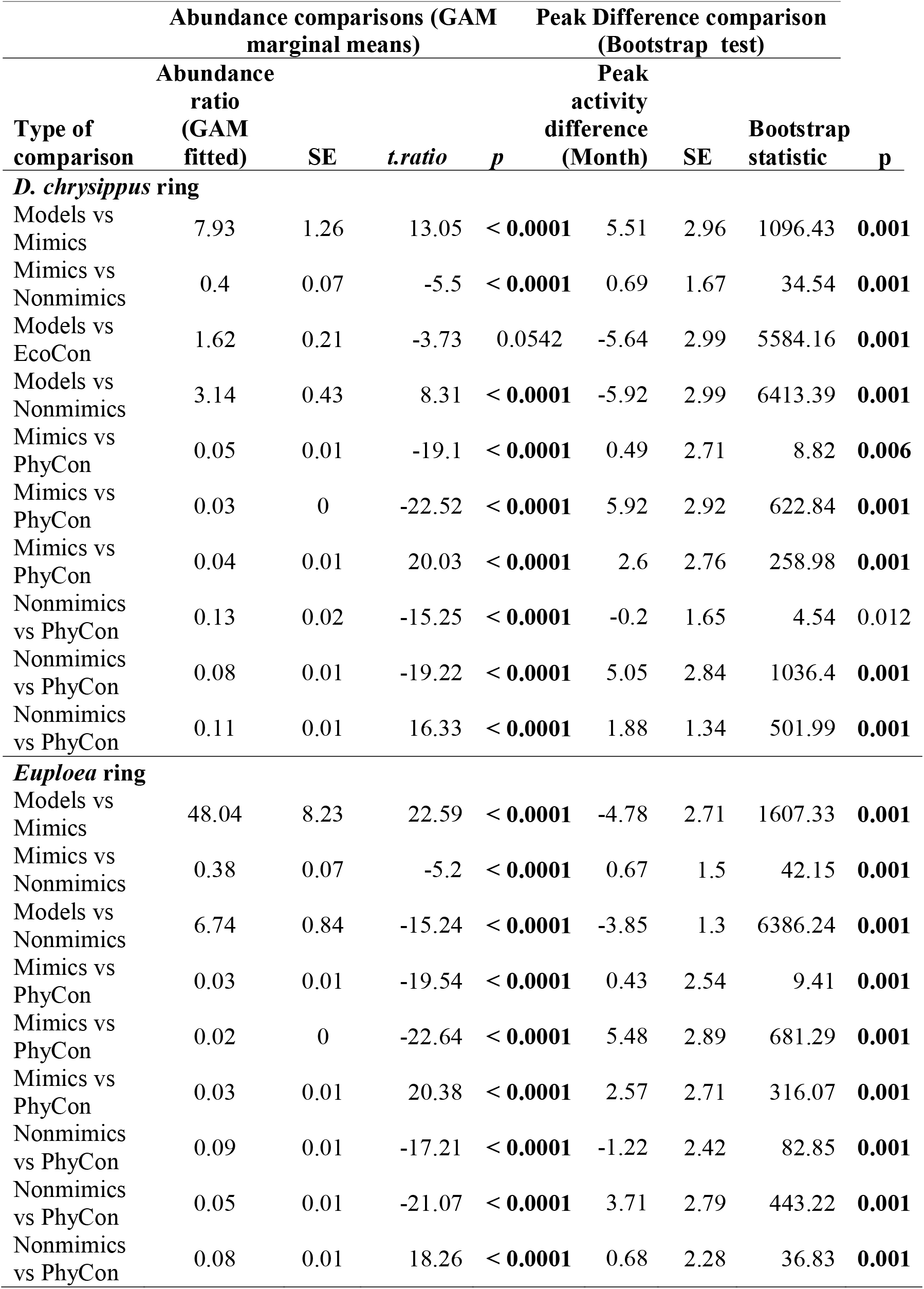

**Table 3C.** Comparison of seasonal population abundance across functional categories of mimetic butterfly communities and their phylogenetic and ecological contrasts. Abundance ratio obtained through marginal means of fitted GAM models. Peak activity difference is obtained by mean of pairwise differences of bootstrapped replicate. Functional groups are shortened for better readability: Models: Batesian models, Mimics: Batesian mimics, Nonmimics: Non-mimetic forms of Batesian mimics, EcoCon: Ecological contrasts of Batesian models, PhyCon: Phylogenetics contrasts of Batesian mimics.

| Type of comparison | Abundance comparisons (GAM marginal means) |  |  |  | Peak Difference comparison (Bootstrap test) |  |  |  |
| --- | --- | --- | --- | --- | --- | --- | --- | --- |
|  | Abundance ratio (GAM fitted) | SE | *t.ratio* | *p* | Peak activity difference (Month) | SE | Bootstrap statistic | *p* |
| ***Tirumala* ring** |  |  |  |  |  |  |  |  |
| Models vs Mimics | 145.47 | 46.42 | -15.61 | < **0.0001** | -3.48 | 2.2 | 763.51 | **0.001** |
| Mimics vs Nonmimics | 0.05 | 0.02 | -9.17 | < **0.0001** | -1.19 | 2.8 | 99.73 | **0.001** |
| Models vs EcoCon | 8.24 | 1.16 | -14.94 | < **0.0001** | 4.43 | 3.07 | 8724.86 | **0.001** |
| Models vs Nonmimics | 7.35 | 1.03 | -14.24 | < **0.0001** | -4.64 | 0.49 | 46441.41 | **0.001** |
| Mimics vs PhyCon | 0.11 | 0.04 | 6.46 | < **0.0001** | 2.77 | 2.14 | 483.77 | **0.001** |
| Mimics vs PhyCon | 0.21 | 0.07 | 4.53 | **0.0023** | 5.79 | 3.02 | 1519.97 | **0.001** |
| Nonmimics vs PhyCon | 2.28 | 0.39 | -4.75 | **0.0008** | 3.93 | 0.44 | 20216.37 | **0.001** |
| Nonmimics vs PhyCon | 4.1 | 0.81 | -7.18 | < **0.0001** | -5.02 | 0.34 | 86750.93 | **0.001** |
| ***D. genutia* ring** |  |  |  |  |  |  |  |  |
| Models vs Mimics | 9.2 | 1.76 | 11.63 | < **0.0001** | -1.89 | 1.17 | 1254.5 | **0.001** |
| Mimics vs Nonmimics | 0.28 | 0.06 | -6.36 | < **0.0001** | 0.12 | 1.15 | 5.02 | 0.019 |
| Models vs Nonmimics | 2.55 | 0.38 | 6.28 | < **0.0001** | -1.78 | 0.77 | 2446.23 | **0.001** |
| Mimics vs PhyCon | 0.19 | 0.04 | -8.41 | < **0.0001** | -0.78 | 1.17 | 238.17 | **0.001** |
| Mimics vs PhyCon | 0.13 | 0.02 | -10.78 | < **0.0001** | -3.03 | 2.3 | 1756.64 | **0.001** |
| Mimics vs PhyCon | 0.01 | 0 | -24.15 | < **0.0001** | -2.24 | 1.16 | 1400.47 | **0.001** |
| Nonmimics vs PhyCon | 0.68 | 0.11 | -2.42 | 0.83 | -0.91 | 0.73 | 742.45 | **0.001** |
| Nonmimics vs PhyCon | 0.45 | 0.07 | -5.25 | < **0.0001** | -3.13 | 1.09 | 2889.48 | **0.001** |
| Nonmimics vs PhyCon | 0.04 | 0.01 | -22.5 | < **0.0001** | -2.34 | 0.96 | 2862.19 | **0.001** |

### b. Population sizes of mimetic and non-mimetic forms of mimetic species are decoupled

Population sizes differed markedly between non-mimetic and Batesian mimetic forms of the same species across all mimetic communities, with non-mimetic males and females consistently exhibiting significantly higher abundances compared with Batesian mimetic females (Table 3, Fig. 3). This pattern was particularly striking considering that mimetic females and non-mimetic males/females of the same species use the same larval host plants (Fig. S4) and are exposed to the same climatic conditions in developmental and adult stages. This suggested that environmental factors alone did not account for the observed differences in their seasonal population abundances, their trajectories appearing to diverge substantially despite their ecological and developmental overlap. This disparity between the abundances of mimetic and non-mimetic forms of the same species was further reflected in the strongly skewed ratios observed in the field, with non-mimetic forms (i.e., non-mimetic males and female forms) greatly outnumbering mimetic forms (all females at our study site) of the same species (mimetic form/non-mimetic form ratio observed in the field = 0.23, SE=0.02; Table 3, 4, Fig. 2). In contrast, the population dynamics of non-mimetic forms of the mimetic species closely resembled those of their non-mimetic phylogenetic contrasts, with comparable abundance levels (Non-mimetic/Phylogenetic contrasts ratio = 1.22, SE=0.1; Fig. 2, Table 3, 4). Together, these patterns indicated that while seasonal population abundances of non-mimetic butterflies matched broader patterns of seasonal abundance of aposematic and non-mimetic species (ecological and phylogenetic contrasts), Batesian mimetic butterflies exhibited distinctive population dynamics, potentially shaped by mimicry relationships with their Batesian models rather than by climatic factors. The potential impacts of mimicry dynamics on the seasonal population dynamics of mimetic butterflies are further tested in the following sections based on predictions of the mimicry theory, specifically, prey phenology under effects of species interactions, seasonal predation risks, and negative frequency-dependent selection (Kunte et al. 2021; Fig. 1).

### c. Population peaks of Batesian models overlap with the seasonal peak of naïve predators

The breeding season of most insectivorous avian predators in the study area span from March to July (Fig. S6), with the latter part of this period coinciding with the fledging of juveniles that begin foraging independently. Consequently, this window (May–July) may be considered a peak phase of naïve predator abundance, when inexperienced individuals are likely to exert distinct selective pressures on prey communities. Notably, populations of Batesian model species across nearly all mimicry rings—except the *Pachliopta hector* ring— reach their peak during this period of heightened naïve predator activity (Fig. 3, S6, Table 5), suggesting a temporal alignment that may enhance predator learning dynamics.

**Table 5.** Average and harmonic mean of abundances across the year for functional categories in mimetic communities. Confidence interval for peak activity calculated by bootstrapping.

| **Mimicry ring** | **Functional category** | **Average abundance across year** | **Harmonic mean abundance across year** | **Peak Activity (Month [95% CI])** |
| --- | --- | --- | --- | --- |
| ***Pach. aristolochiae* ring** | Batesian models | 6.27 | 4.33 | 7.95 [7.3–10.3] |
|  | Batesian mimics | 0.9 | 0.73 | 6 [4.4–10.1] |
|  | Non-mimetic forms | 7.04 | 2.56 | 6.9 [6–8.7] |
| ***Pach. hector* ring** | Batesian models | 3.05 | 2.96 | 2.7 [10.3–7.2] |
|  | Batesian mimics | 0.9 | 0.74 | 7.5 [5.8–10.4] |
|  | Non-mimetic forms | 7.04 | 2.56 | 6.9 [6–8.7] |
| ***D. chrysippus* ring** | Batesian models | 6.39 | 5.85 | 5.4 [2.7–6.8] |
|  | Batesian mimics | 0.8 | 0.63 | 10.5 [9.8–3.55] |
|  | Non-mimetic forms | 2.07 | 1.5 | 10.7 [10.5–12.9] |
| ***Euploea* ring** | Batesian models | 33.17 | 21.6 | 6.8 [5.6–8.4] |
|  | Batesian mimics | 0.56 | 0.41 | 10.6 [9.9–3] |
|  | Non-mimetic forms | 1.39 | 1.3 | 10.3 [5.8–10.9] |
| ***Tirumala* ring** | Batesian models | 24.84 | 7.56 | 5.4 [4.6475–6.2] |
|  | Batesian mimics | 0.09 | 0.08 | 9.5 [4.1–3] |
|  | Non-mimetic forms | 1.94 | 1.32 | 10.1 [9.45–10.4] |
| ***D. genutia* ring** | Batesian models | 3.88 | 3.17 | 7.4 [6.3–8.7] |
|  | Batesian mimics | 0.4 | 0.32 | 9.5 [7.3–10.8] |
|  | Non-mimetic forms | 1.77 | 1.05 | 9.2 [8–10.1] |

On the other hand, ecological contrasts of the Batesian models (i.e., aposematic species existing without any Batesian mimics) exhibited significantly different population dynamics, with pronounced temporal mismatches in peak activity. Specifically, Batesian models peaked substantially earlier than their ecological contrasts, with a significant difference in peak timing (mean difference = 2.94, SE = 2.23; Tables 3–4; Fig. 4). Additionally, Batesian models were far more abundant than their ecological contrasts, as reflected in a markedly elevated abundance ratio (Batesian models/ecological contrasts ratio = 18.46, SE=0.02; Table 3–4; Fig. 3). Together, these patterns highlight strong temporal and numerical advantages to Batesian models relative to comparable non-model aposematic species, particularly during periods of high naïve predator pressure.

**Figure 4.**
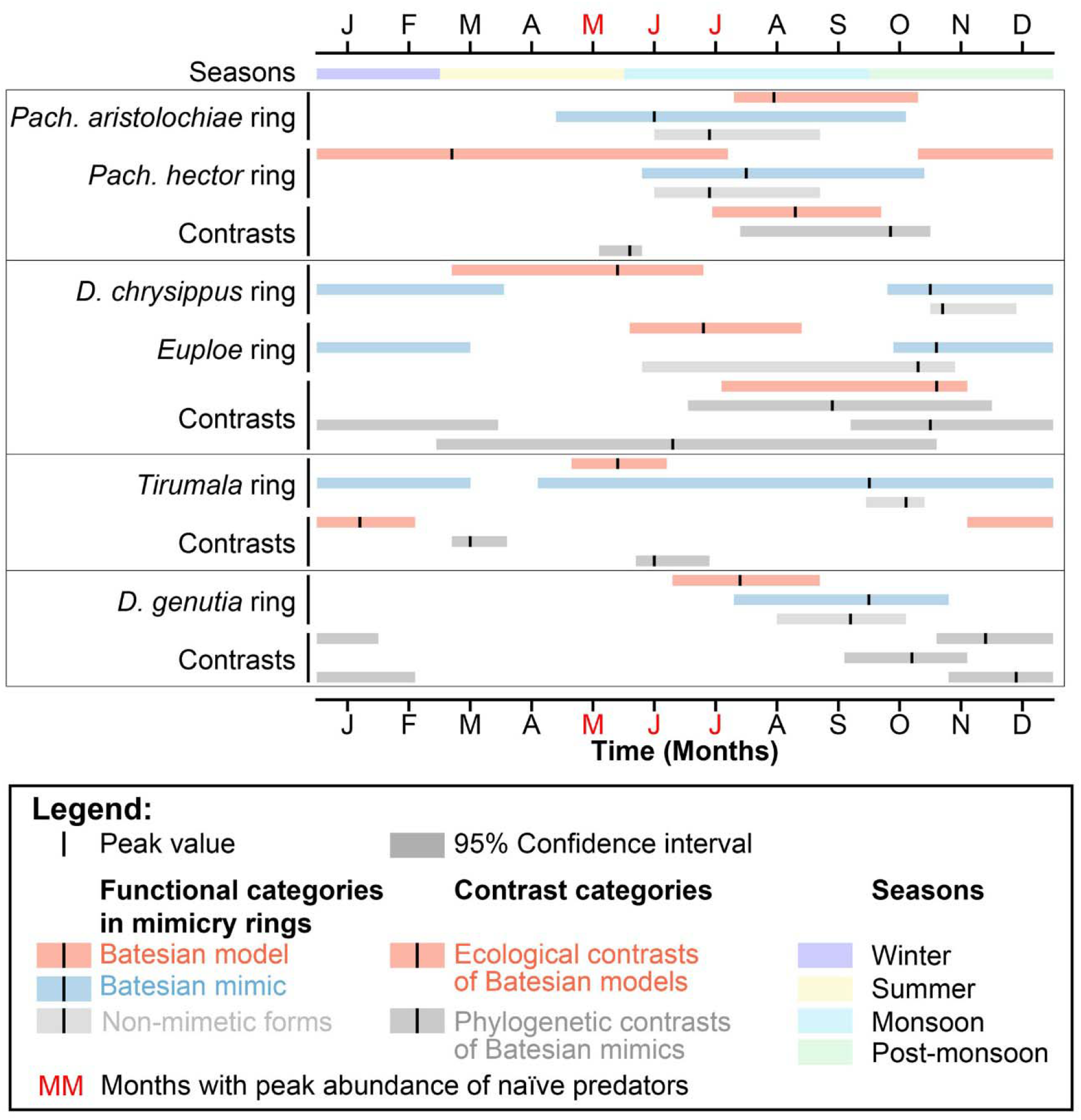
Seasonal population peaks of Batesian model overlap with the season of peak naïve avian predators, whereas seasonal population peaks of Batesian mimics are positively phase-shifted compared to their Batesian models. Activity peaks, with 95% confidence intervals, obtained from bootstrapped values of GAM fitted abundance curves, are plotted.

**Table 4.**
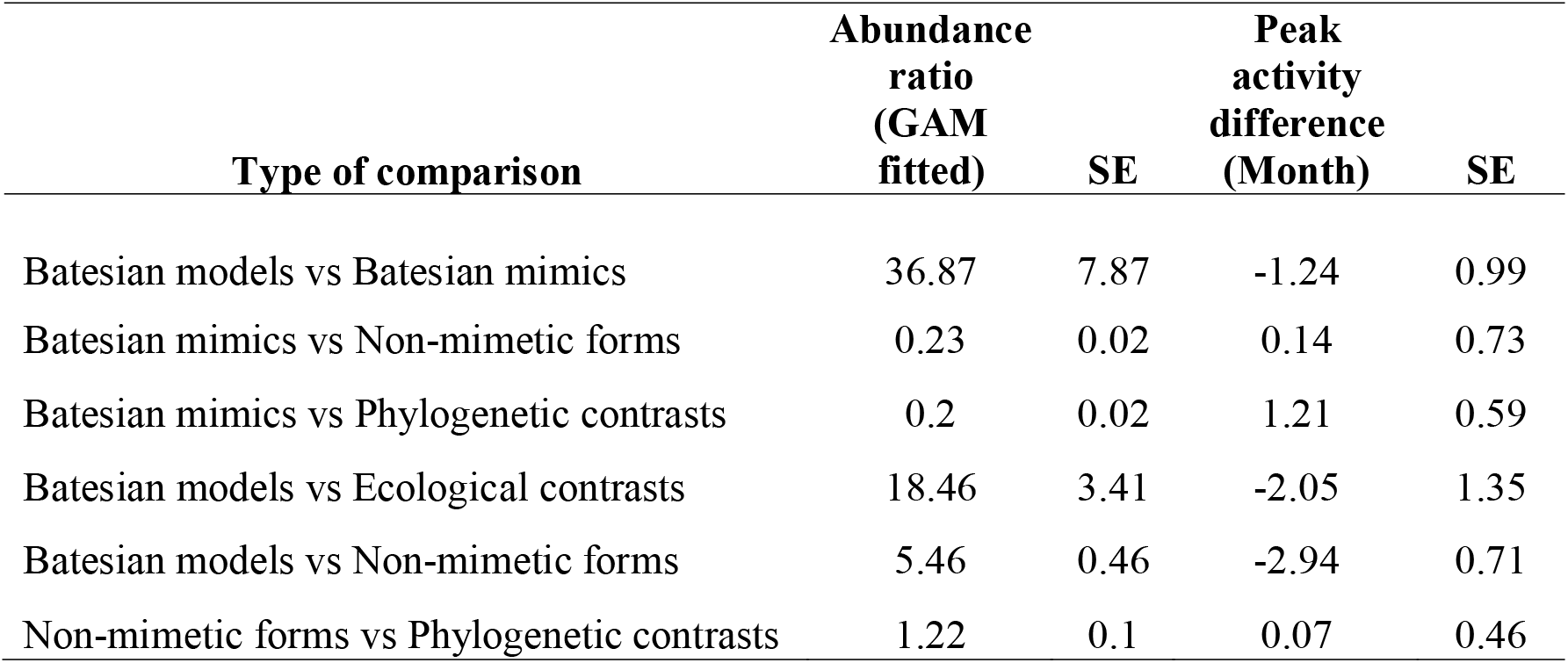
Mean comparison across functional categories. Abundance ratio obtained through marginal means of fitted GAM models. Peak activity difference is obtained by mean of pairwise differences of bootstrapped replicate with peak of first species minus second. Thus, negative value indicates first species in the comparison peaking early.

### d. Batesian models were more abundant, with Batesian mimics peaking later

Across all mimicry rings, Batesian models were substantially more abundant than their corresponding Batesian mimics (Model/Mimic ratio = 36.87, SE 7.87, Tables 3–4). In addition, non-mimetic forms within mimetic species were consistently more abundant than both their mimetic counterparts and the aposematic models that they resembled. This hierarchical pattern in abundance was not limited to standardized surveys but was also evident in independent opportunistic records from the citizen science dataset (Fig. S3), indicating that it is robust across sampling approaches.

Temporal dynamics, especially population peaks, further distinguished Batesian models from mimics. Batesian models and their mimics exhibited slight differences in peak activity, with mimics reaching peak abundance after the models (peak difference=1.24, SE=3.93; Tables 3–4, Figs. 3–4). Importantly, the peak activity of Batesian models closely coincided with the period of highest naïve predator abundance, whereas mimics showed a consistent positive phase shift relative to the models and peak naïve predator season (except in the case of *Pachliopta aristolochiae* mimicry ring). This temporal offset suggests that models primarily interact with naïve predators, facilitating predator learning before the population peaks of Batesian mimics. Thus, Batesian mimic prey populations may experience relatively relaxed selection from a predator community that experienced by the time of the population peaks of the mimetic prey.

## DISCUSSION

Climate emerges as a key driver of butterfly population dynamics, with the rainy season consistently associated with higher abundances across the annual cycle (Bonebrake et al. 2010; Checa et al. 2019; Colom et al. 2022). This effect is both direct and indirect: favourable climatic conditions not only enhance survival and activity but also influence the availability and quality of larval host plants, which are tightly linked to rainfall and temperature regimes (Larsen et al. 2022; Navarro-Cano et al. 2015). Consequently, optimal climatic conditions translate into improved resource availability, supporting larger populations. Consistent with this, population abundance of non-mimetic forms and the phylogenetic contrasts of Batesian mimics showed strong associations with climatic variables, with a substantial proportion of variation in their population sizes explained by temperature, precipitation, and relative humidity. In contrast, Batesian mimics exhibited a markedly weaker relationship with these climatic drivers, with relatively low variance in abundance attributable to climatic parameters. Together, these results suggest that while climatic factors strongly structure the population dynamics of non-mimetic, aposematic and phylogenetically related taxa, they play a comparatively limited role in shaping the population dynamics of Batesian mimics, pointing to the importance of additional ecological or evolutionary processes in these systems (Hassall, Billington, and Sherratt 2019).

Population dynamics of Batesian models appear to be closely aligned with periods of increased naïve predation pressure, with their abundance peaks typically overlapping with or closely following peaks in naïve predator activity. In (sub)tropical systems, overall predation pressure remains relatively constant throughout the year because predator communities are continuously active (Repetto et al. 2024; Sam et al. 2026); however, the composition of that pressure changes seasonally. In particular, the post-breeding period—typically during resource-rich rainy season—leads to a surge of inexperienced, visually-hunting naïve predators such as birds entering the foraging population. This influx of naïve predators likely imposes disproportionately high predation pressure on conspicuous aposematic species, as these predators have not yet learned to associate warning signals with unprofitability. By extension, Batesian mimics may also experience increased risk during this period. Under such conditions, higher abundances of Batesian models can be advantageous for both Batesian models and mimics, as they facilitate more rapid predator learning and reinforce avoidance behaviour without hindering learning by naïve predators with a mix of edible Batesian mimics. Thus, relatively lower abundances of mimics during this period of peak naïve predator abundance may stabilize the system by maintaining the reliability of warning signals during predator learning. These dynamics are reflected in the observed phenological patterns, where Batesian mimics exhibited a consistent positive phase shift relative to their models, likely constrained by the need to track periods when predator communities are more experienced in avoiding unprofitable prey. This temporal offset may, in turn, contribute to divergent population trajectories between non-mimetic males and Batesian mimetic females within species. While mimetic females are subject to constraints imposed by mimicry dynamics and predator learning, non-mimetic males are not bound by these interactions and instead show population patterns more closely associated with climatic drivers (Tsurui Sato et al. 2019).

Population dynamics of Batesian mimics are likely closely linked to those of their corresponding models, with the abundance of mimetic females effectively constrained by frequency-dependent selection relative to model abundance, imposed by the negative frequency-dependent selection imposed by Batesian mimicry. Because the success of mimicry depends on maintaining a low ratio of mimics to models, increases in mimic frequency reduce signal reliability and lead to higher predation risk, thereby suppressing mimic abundance (Kunte et al. 2021). One way this constraint may be mitigated is through the compartmentalization of mimicry within a specialized subset of the population, such as sex-limited and/or polymorphic mimicry (Basu et al. 2023; Kunte 2009; Kunte et al. 2021). By restricting the mimetic phenotype to a relatively rare form, the species can limit the overall “mimetic load” imposed on the system, effectively avoiding the stricter bounds of frequency-dependent selection that would apply if all individuals were mimetic (Kunte 2008, 2009). This division allows the species to balance the benefits of mimicry with the costs of signal dilution.

At the same time, Batesian mimetic females—being directly exposed to predation during oviposition and other reproductive activities—may derive substantial fitness advantages from resembling unpalatable models. Enhanced survival of egg-laying females can translate into higher reproductive output, thereby benefiting population persistence despite constraints on mimic frequency (Kunte 2008, 2009). Together, these dynamics suggest that sex-limited mimicry may represent an adaptive strategy that reconciles the opposing pressures of predator-mediated selection and frequency dependence (Kunte 2009; Kunte et al. 2021).

The frequency of mimics relative to models, calculated as mimic abundance / (Model + Mimic abundance), was estimated at 0.10±0.07. This value is substantially lower than those reported in previous studies. For instance, laboratory experiments using artificial prey and starling predators suggest that effective predator learning and avoidance of mimics occur only when mimic frequencies remain below approximately 0.3–0.6 (Brower 1960). However, such estimates likely overstate thresholds applicable to natural systems, as they typically assume highly toxic models and near-perfect mimicry. Long-term field studies provide more realistic benchmarks. A 15-year study reported mimic frequencies of 0.3±0.09 in a Batesian mimicry system and 0.35±0.1 in a female-limited polymorphic system (Long, Edwards, and Shapiro 2015). In contrast, other work has documented substantial geographic variation, with frequencies ranging from 0.31 to 0.99 (Prudic et al. 2019). These higher and more variable estimates likely reflect systems in which mimics possess some degree of unpalatability or function as Müllerian co-models, thereby relaxing strict frequency-dependent constraints by balancing model abundance against the physiological costs of toxin sequestration. In comparison, our results indicate that (sub)tropical mimetic communities with continuously breeding populations operate under much stricter frequency-dependent limits. We propose that the harmonic mean of model abundance—by weighting periods of low abundance more heavily—provides a more appropriate measure of the effective constraint imposed by predator-mediated selection (Table 5, Fig. 3) and should be used for mimic frequency estimation. This approach is analogous to use of harmonic mean in estimating effective population size in temporally fluctuating populations, where rare but critical low-frequency phases disproportionately influence effective population size and resultant long-term adaptive dynamics (Fedorca et al. 2024; Waples 2025).

The consistently higher abundance of Batesian models relative to their ecological contrasts suggests two alternative evolutionary interpretations. First, models may need to maintain larger population sizes to buffer the parasitic load imposed by Batesian mimics, ensuring the stability of the warning signal despite exploitation. Alternatively, the prior existence of an abundant aposematic species may be a prerequisite for the evolution of mimicry, providing a sufficiently strong and reliable signal for mimetic selection. We consider the second explanation more plausible. For example, one of the ecological contrasts (*Delias eucharis*) participates in a mimicry ring with a Western Ghats endemic species that is locally abundant within the mimic’s range, indicating that high model abundance should precede and facilitate the evolution of mimicry (Kunte et al. 2021). Under this view, Batesian mimicry arises in response to pre-existing, abundant aposematic models, rather than driving increases in model abundance (Kunte et al. 2021). Once established, however, the persistence and structure of these mimicry systems are likely governed by a combination of seasonal climatic variation, predator-mediated selection, and strict frequency-dependent constraints. Together, these forces shape both the temporal dynamics and relative abundances of models and mimics within (sub)tropical communities.

Overall, our study provides insight into the complex population dynamics of a subtropical system characterized by highly fluctuating, continuously breeding prey populations. In general, peak abundances and broader population trajectories are shaped by climate and the availability of host plants, with predation acting as an additional, strategy-dependent regulatory force. While background predation remains relatively constant throughout the year, seasonal surges of naïve predators impose disproportionately high risk on conspicuous aposematic species, which are both highly visible and relatively easy to capture.

Consequently, predation emerges as a key driver of population dynamics in aposematic prey and, by extension, in mimetic species constrained by frequency-dependent selection. This predator-mediated pressure, operating in conjunction with climatic and resource-driven processes, likely underpins the temporal structure and abundance patterns observed across mimicry rings. Frequency-dependent selection thus emerges as a central force shaping both the evolution and maintenance of mimicry, including the origin and persistence of specialized strategies such as female-limited mimicry (Kunte et al. 2021).

The patterns of seasonal population abundance of functional groups within mimetic butterfly communities and their phylogenetic and ecological contrasts described in this work broadly support predictions of mimicry theory with respect to negative frequency-dependent selection and resultant prey phenology (Kunte et al. 2021) (Fig. 1). Importantly, these patterns are consistent across multiple mimetic communities, suggesting that seasonal abundance patterns and underlying ecological processes apply consistently and broadly in the context of functional groups within ecological communities, irrespective of individual species biology. Thus, it is possible to view community evolution and phenology within community ecological and evolutionary ecological frameworks, with functional groupings underpinning the broad patterns. Such a focus on functional groups is not always apparent in many ecological studies. Our use of suitable phylogenetic and ecological contrasts, and their value in separating broad functional group-level patterns from narrower species-level patterns, should further motivate a broader adoption of the use of and comparison across functional groups and phylogenetic/ecological contrasts in classical community ecological studies.

## Acknowledgments

We thank members of the Bangalore Butterfly Club for assistance in data collection; Ujwala Pawar for some of the trait dataset; and Deepa Agashe for advice on tree models analysis. This work was supported by an NCBS graduate student fellowship to VN, personal funds from RG and NR; and an NCBS research grant to KK.

## Author Contributions

VN curated and analysed data, and prepared initial drafts of figures and the paper; RG and NR collected and compiled butterfly count data; KK conceived, coordinated and supervised the study, provided guidance on larval host plant and predator data, identified ecological and phylogenetic contrasts, and contributed some parts of the manuscript; V.N. and K.K. prepared final figures and final draft of the paper.

---

**Fig. S1.**
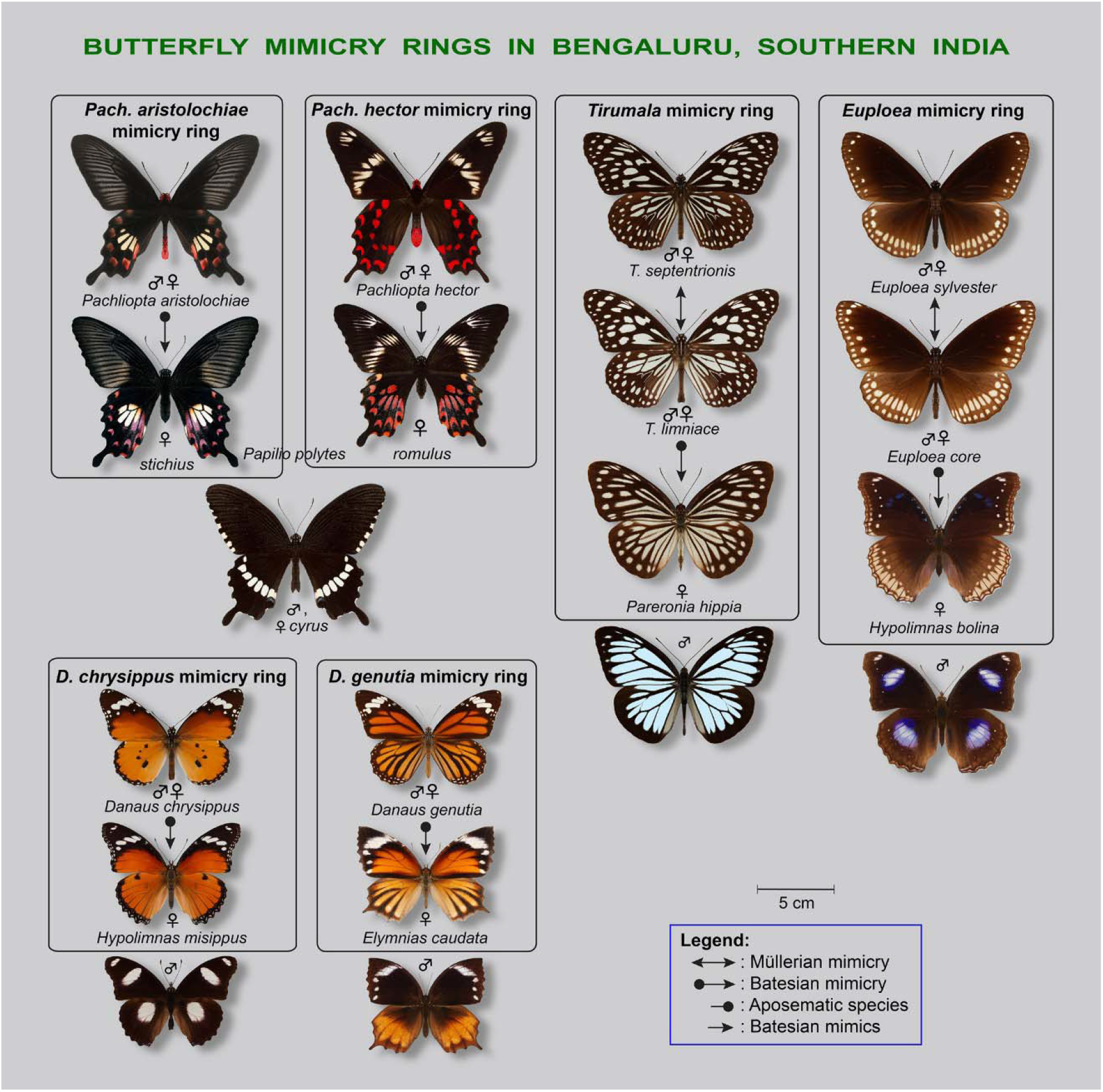
Butterfly mimicry rings present at the study site in Bengaluru, southern India. Monomorphic species are represented by a single image, while each form in sexually dimorphic and polymorphic species is shown separately. Boxes enclose the core components of each mimetic interaction, with non-mimetic forms depicted outside the boxes.

**Fig. S2.**
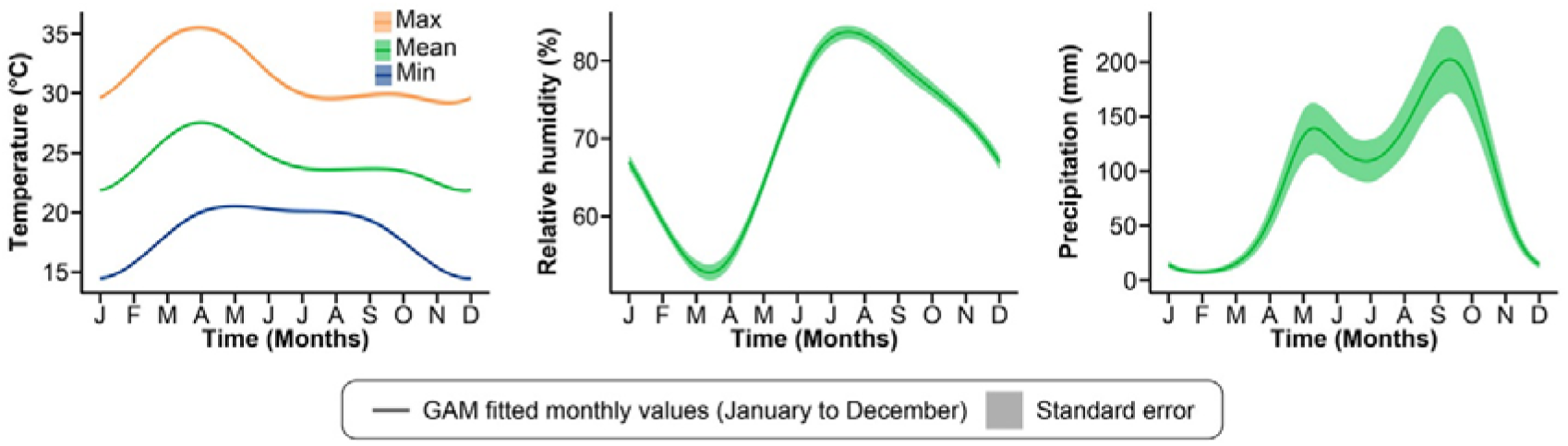
Fitted GAM with climate data showing yearly pattern for each climatic parameter.

**Fig. S3.**
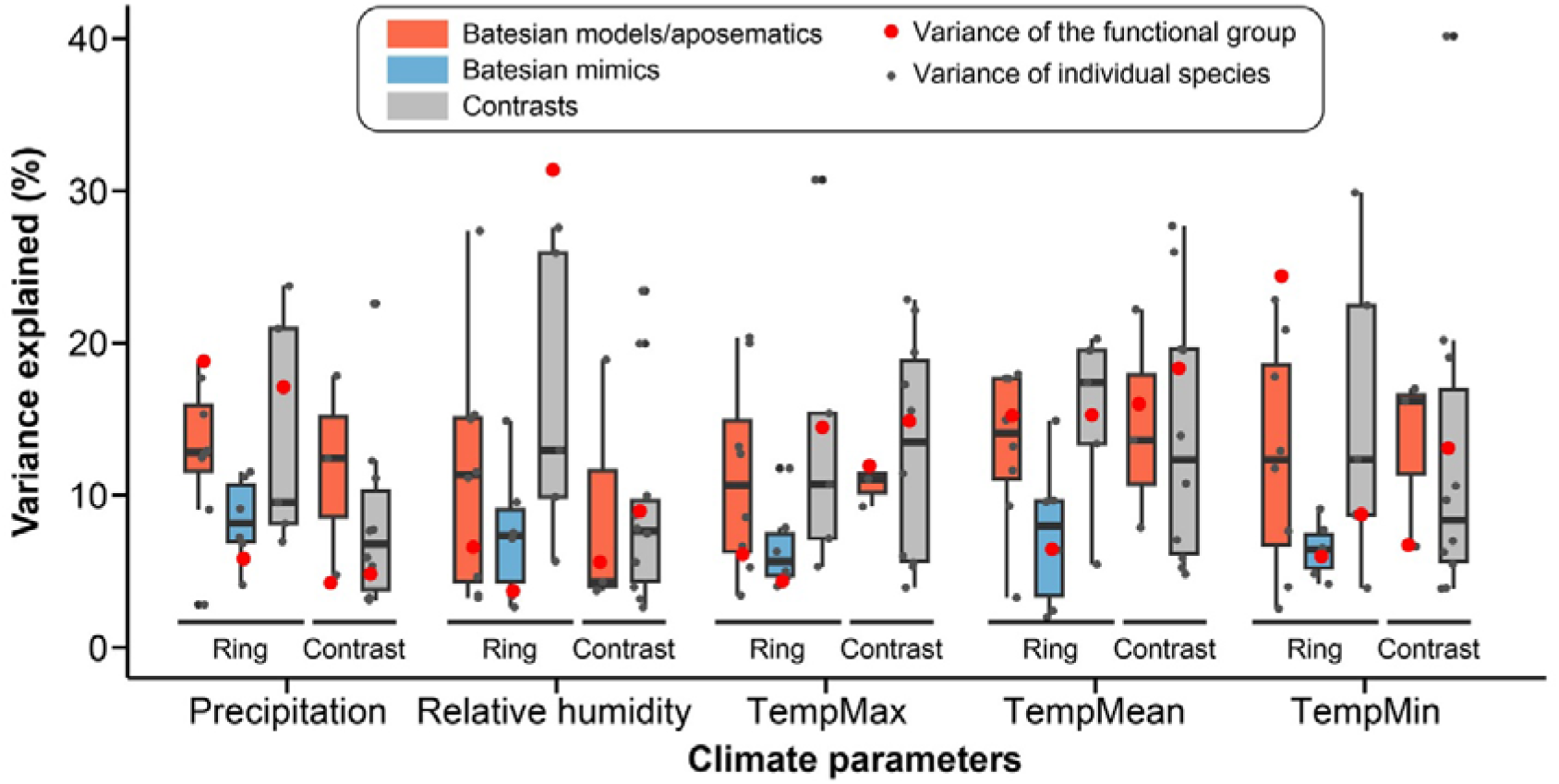
Comparison of variance explained by first split for the functional group compared to individual species part of each group.

**Fig. S4.**
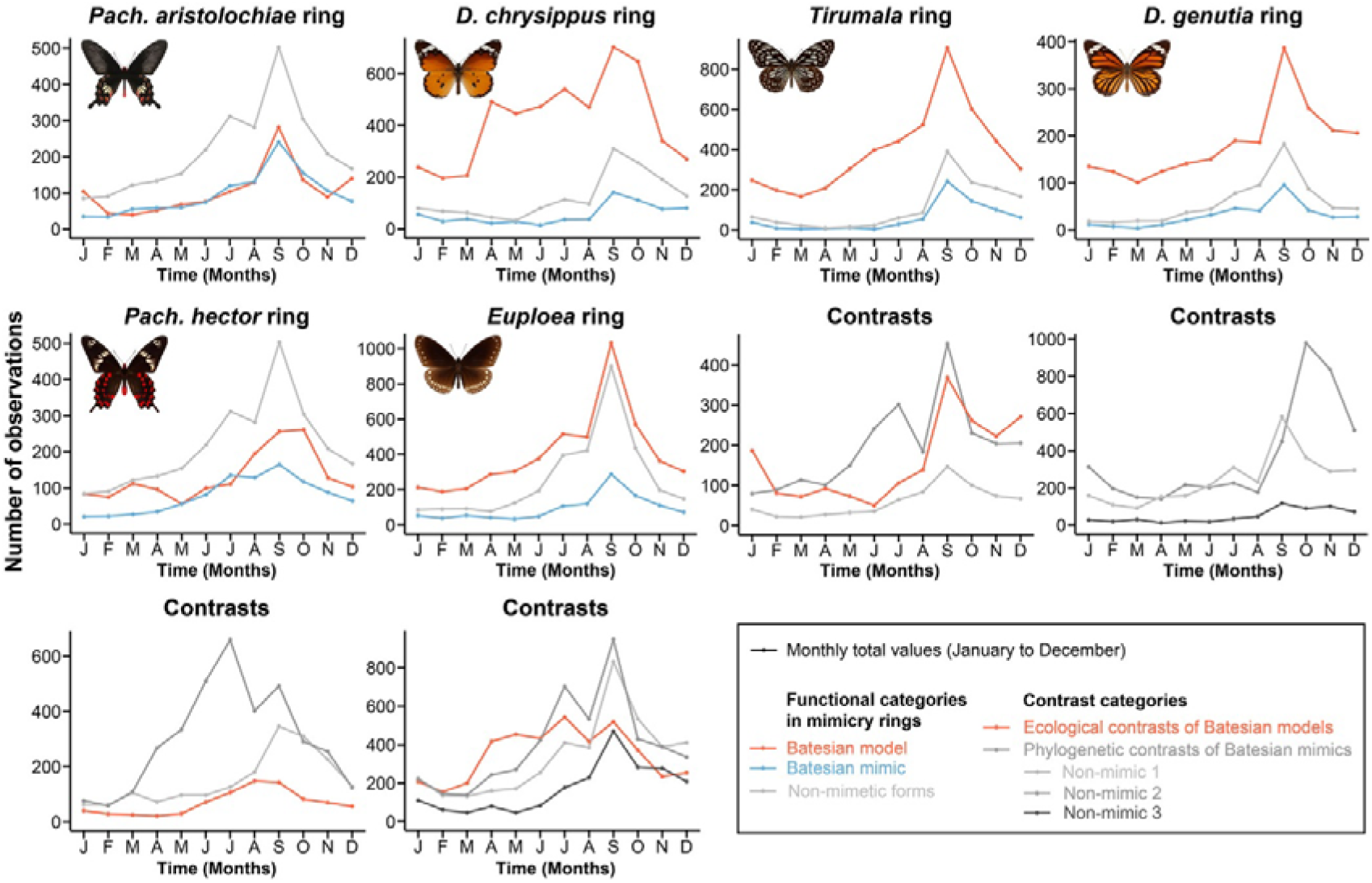
Number of observations per month from citizen science platforms for species in various mimicry rings and their phylogenetic and ecological contrasts are shown. For complete species legend refer Fig. S6.

**Fig. S5a.**
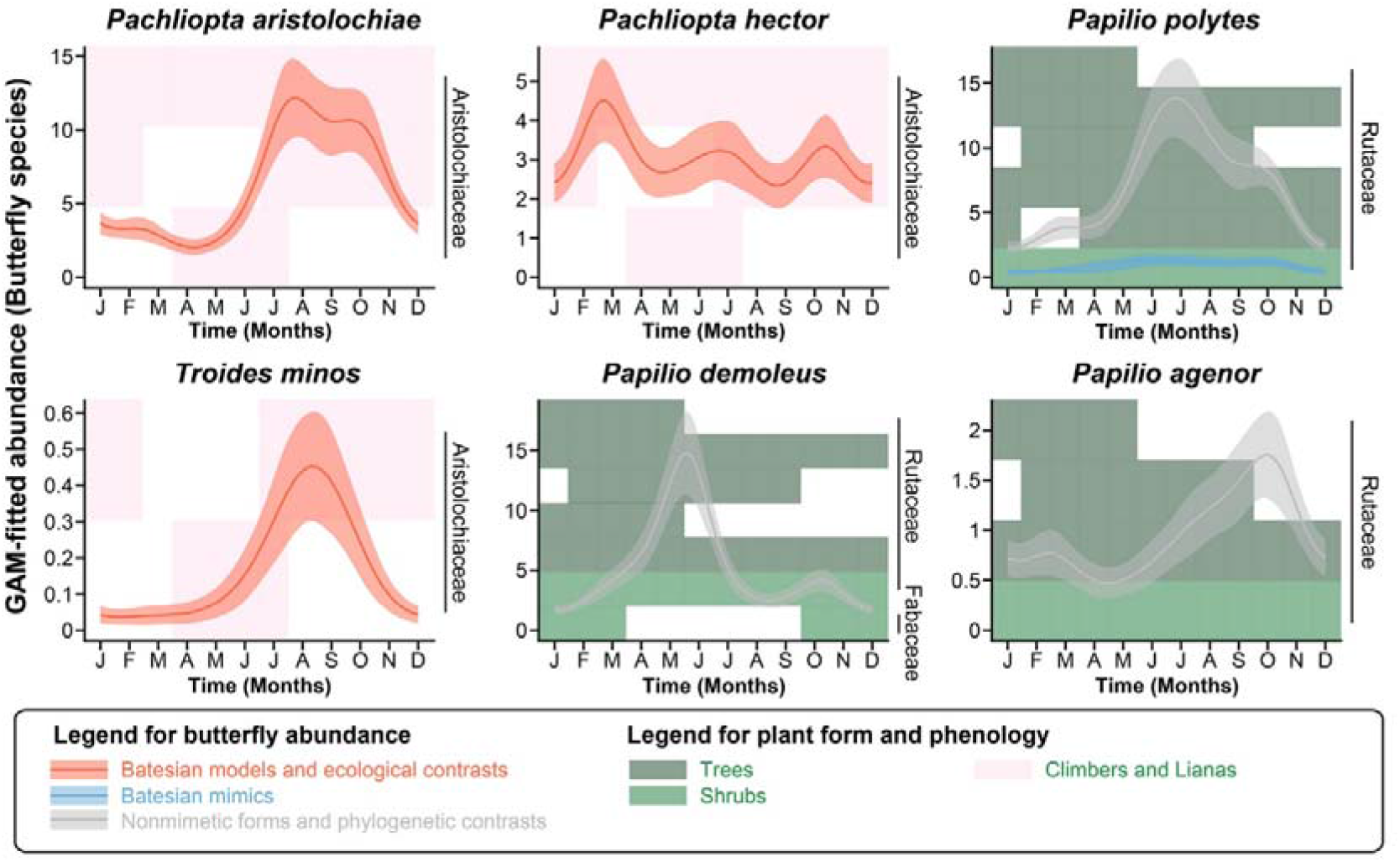
Host plant phenology for species in *Pachliopta aristolochiae* and *Pachliopta hector* mimicry rings. Filled rectangles represent host plant in flowering/fruiting stage. Host plant species are colour coded for their habit with host plant family indicated on the right.

**Fig. S5b.**
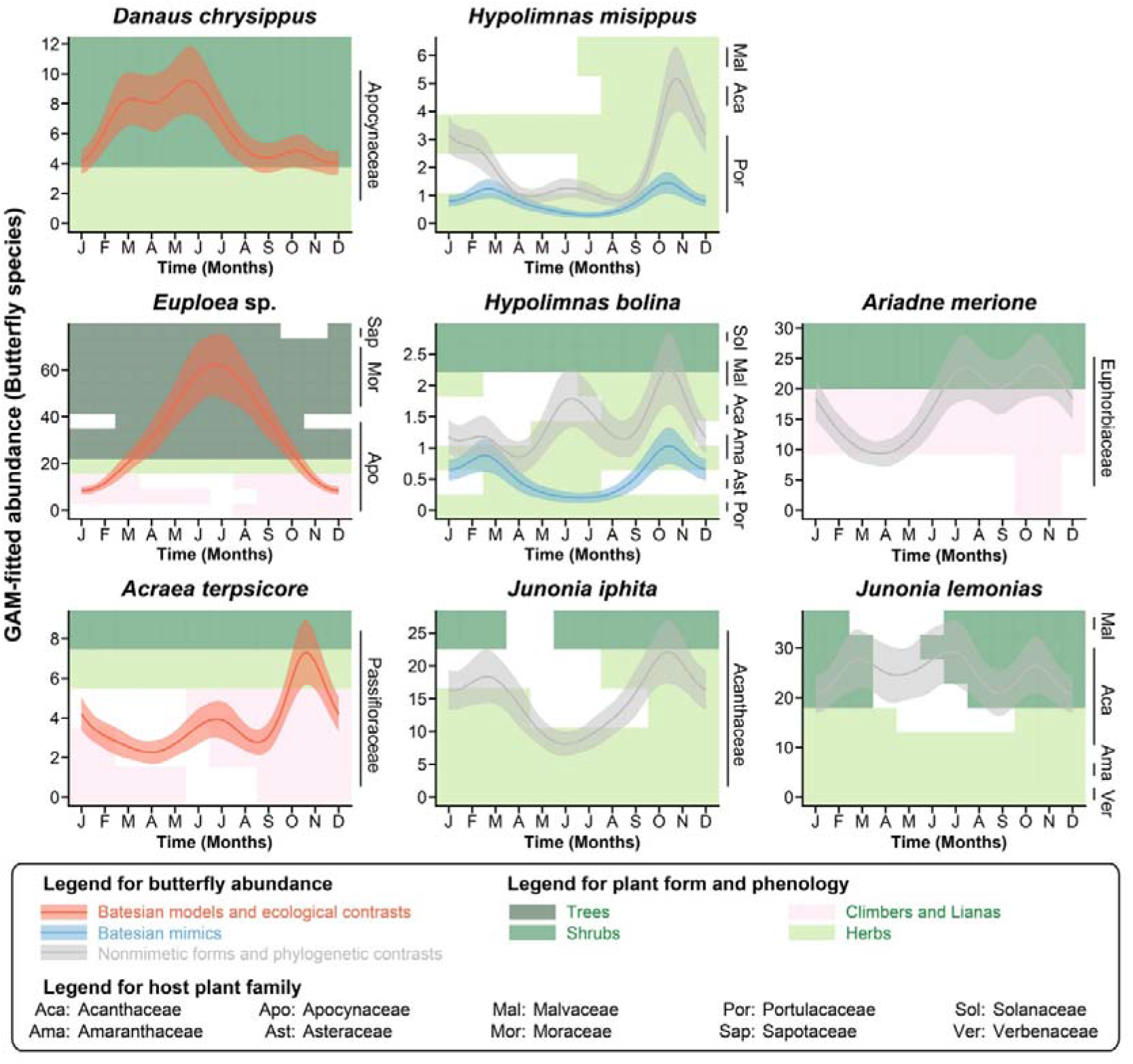
Host plant phenology for species in *Danaus chrysippus* and *Euploea* mimicry rings. Filled rectangles represent host plant in flowering/fruiting stage. Host plant species are colour coded for their habit with host plant family indicated on the right.

**Fig. S5c.**
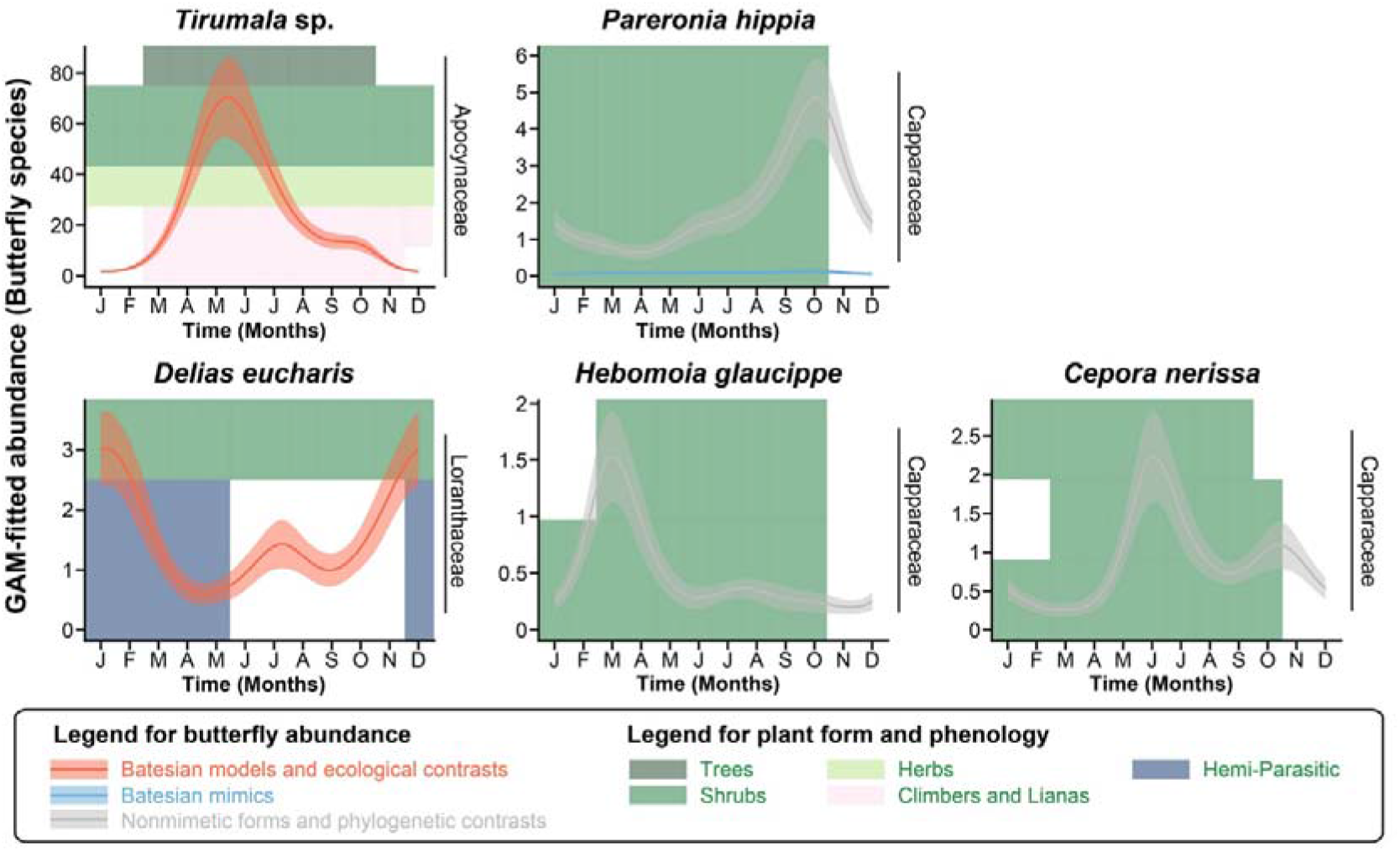
Host plant phenology for species in *Tirumala* mimicry ring. Filled rectangles represent host plant in flowering/fruiting stage. Host plant species are colour coded for their habit with host plant family indicated on the right.

**Fig. S5d.**
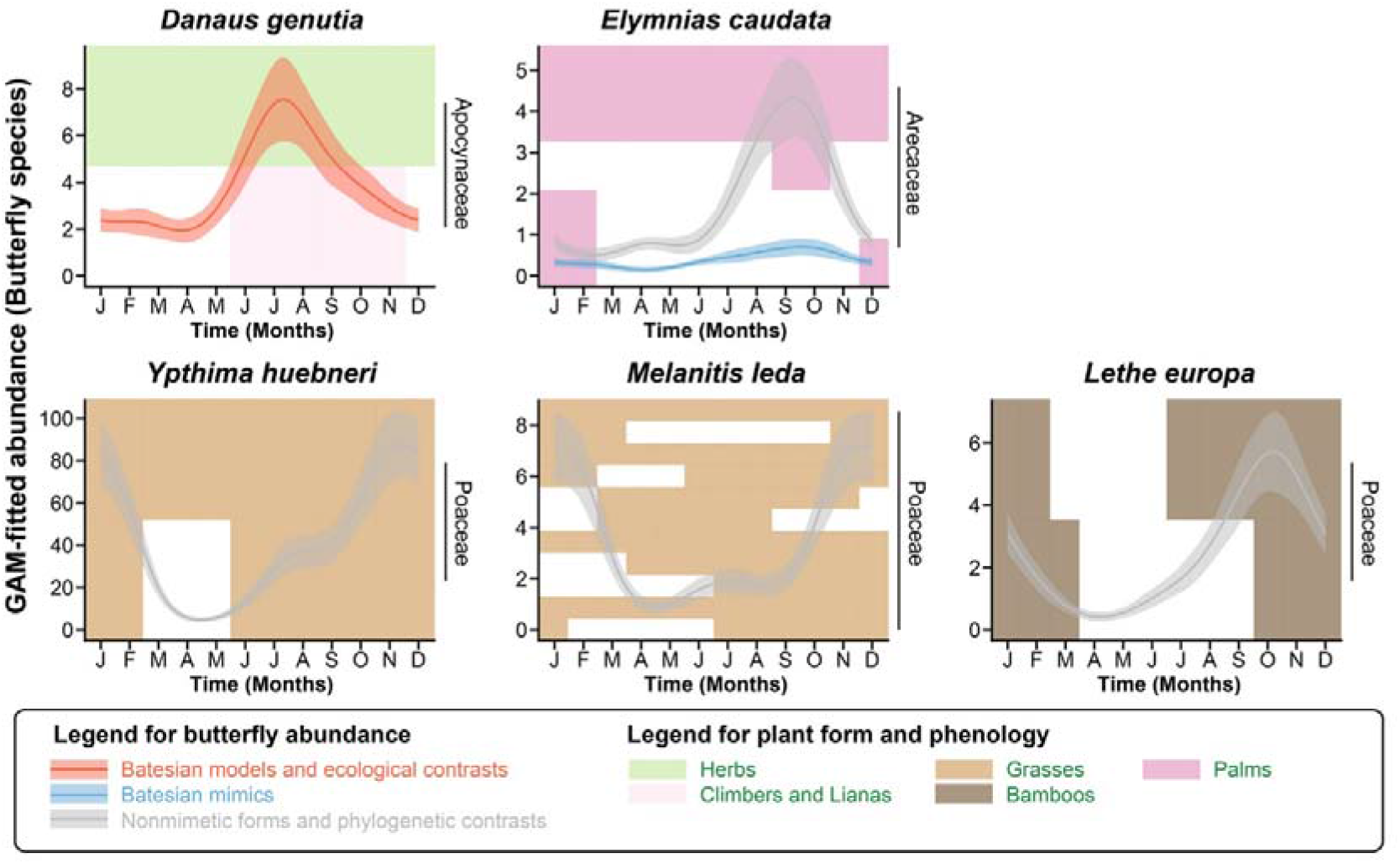
Host plant phenology for species in *Danuas genutia* mimicry ring. Filled rectangles represent host plant in flowering/fruiting stage. Host plant species are colour coded for their habit with host plant family indicated on the right.

**Fig. S6.**
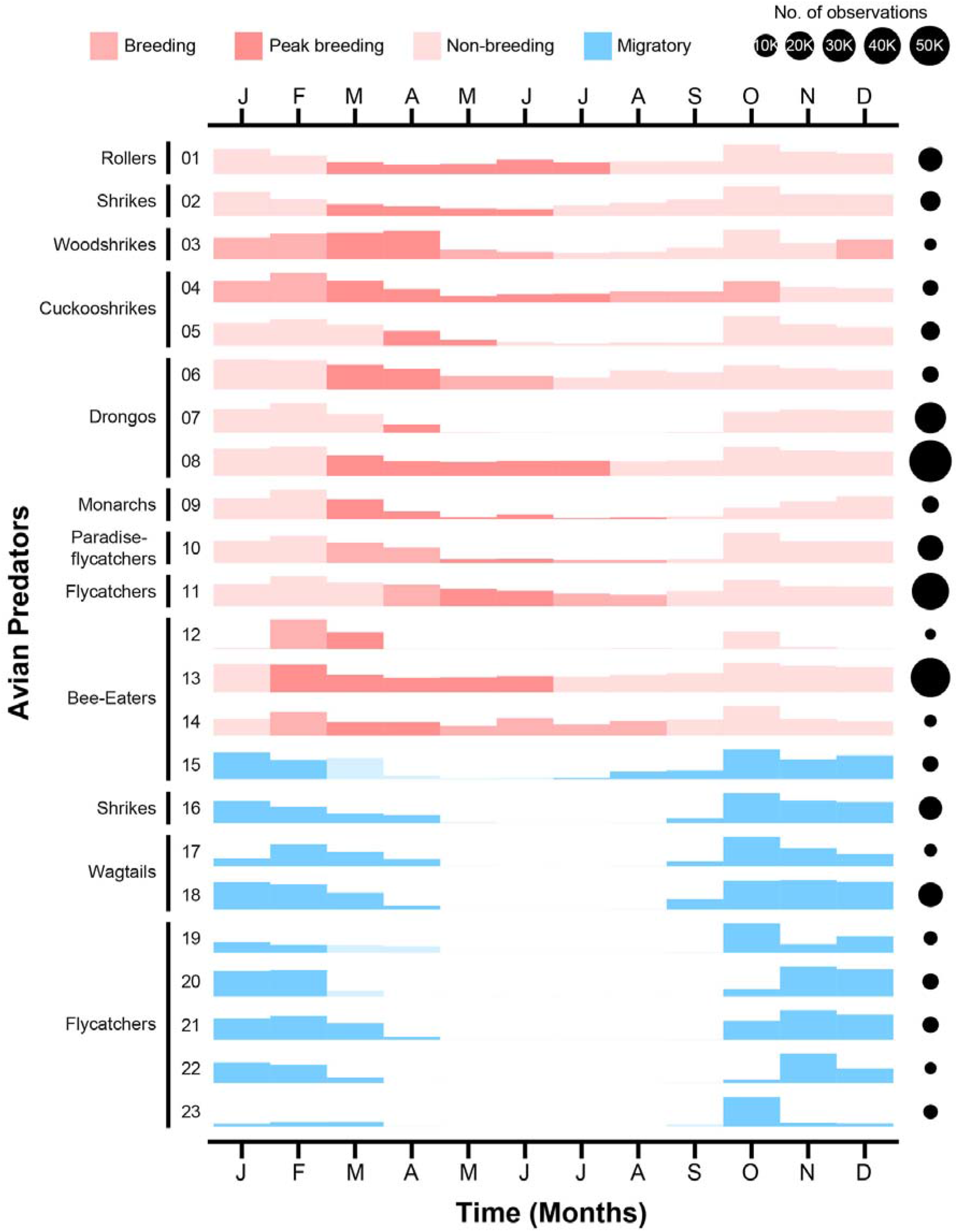
Bird phenology for hawking insectivorous birds which are potential predators of adult butterflies. Height of each bar shows normalized number of observations per month. Circles on the right shows total number of observations for the bird from Bengaluru area.

## References

Ali, S. and S. D. Ripley. 1983. Handbook of the Birds of India and Pakistan, Together with Those of Bangladesh, Nepal, Bhutan and Sri Lanka (Compact Edition). Delhi. Oxford University Press. pp. 737.

Attiwilli, S., N. Ravikanthachari, and K. Kunte. 2024. A comparison between time constrained counts and line transects as methods to estimate butterfly diversity and monitor populations in tropical habitats. Insect Conservation and Diversity, 17:88– 101.

Basu, D. N., V. Bhaumik, and K. Kunte. 2023. The tempo and mode of character evolution in the assembly of mimetic communities. Proceedings of the National Academy of Sciences USA, 120:e2203724120.

Bonebrake, T. C., L. C. Ponisio, C. L. Boggs, and P. R. Ehrlich. 2010. More than just indicators: A review of tropical butterfly ecology and conservation. Biological Conservation, 143:1831–1841.

Both, C., M. Van Asch, R. G. Bijlsma, A. B. Van Den Burg, and M. E. Visser. 2009. Climate change and unequal phenological changes across four trophic levels: constraints or adaptations? Journal of Animal Ecology, 78:73–83.

Breiman, L., J. H. Friedman, R. A. Olshen, and C. J. Stone. 1984. Classification And Regression Trees. Chapman and Hall/CRC.

Brisson, D. 2018. Negative Frequency-Dependent Selection Is Frequently Confounding. Frontiers in Ecology and Evolution, 6.

Brower, J. V. Z. 1960. Experimental studies of mimicry. IV. The reactions of starlings to different proportions of models and mimics. American Naturalist, 94:271–282.

Checa, M. F., E. Levy, J. Rodriguez, and K. Willmott. 2019. Rainfall as a significant contributing factor to butterfly seasonality along a climatic gradient in the neotropics.

Colom, P., M. Ninyerola, X. Pons, A. Traveset, and C. Stefanescu. 2022. Phenological sensitivity and seasonal variability explain climate-driven trends in Mediterranean butterflies. Proceedings of the Royal Society B: Biological Sciences, 289.

Datta, A., S. Banerjee, R. Naniwadekar, Khem Thapa, A. Rathore, Kumar Thapa, T. Brah, T. Nabum, N. Mogar, U. Shukla, S. Sidhu, and N. Borawake. 2025. Patterns of leaf, flower, and fruit phenology and environmental relationships in a seasonal tropical forest in the Indian eastern himalaya. Biotropica, 57.

Edwards, C. B. and E. E. Crone. 2021. Estimating abundance and phenology from transect count data with GLMs. Oikos, 130:1335–1345.

Fedorca, A., J. Mergeay, A. O. Akinyele, T. Albayrak, I. Biebach, A. Brambilla, P. A. Burger, E. Buzan, I. Curik, R. Gargiulo, J. A. Godoy, S. C. González Martínez, C. Grossen, M. Heuertz, S. Hoban, J. Howard McCombe, M. Kachamakova, P. Klinga, V. Köppä, E. Neugebauer, I. Paz Vinas, P. B. Pearman, L. Pérez Sorribes, B. Rinkevich, I. M. Russo, A. Theraroz, N. E. Thomas, M. Westergren, S. Winter, L. Laikre, and A. Kopatz. 2024. Dealing With the Complexity of Effective Population Size in Conservation Practice. Evolutionary Applications, 17.

Fisher, N. I. 1993. Statistical Analysis of Circular Data. Cambridge University Press.

Getman-Pickering, Z. L., G. J. Soltis, S. Shamash, D. S. Gruner, M. R. Weiss, and J. T. Lill. 2023. Periodical cicadas disrupt trophic dynamics through community-level shifts in avian foraging. Science, 382:320–324.

Hassall, C., J. Billington, and T. N. Sherratt. 2019. Climate-induced phenological shifts in a Batesian mimicry complex. Proceedings of the National Academy of Sciences, 116:929– 933.

Hersbach, H., B. Bell, P. Berrisford, G. Biavati, A. Horányi, J. M. Sabater, J. Nicolas, C. Peubey, R. Radu, I. Rozum, and others. 2023. ERA5 hourly data on single levels from 1940 to present. Copernicus Climate Change Service (C3S) Climate Data Store (CDS) [Dataset].

Joshi, J., A. Prakash, and K. Kunte. 2017. Evolutionary assembly of communities in butterfly mimicry rings edited by S. L. Nuismer and J. L. Bronstein. The American Naturalist, 189:E58–E76.

Kishimoto Yamada, K. and T. Itioka. 2015. How much have we learned about seasonality in tropical insect abundance since Wolda (1988)? Entomological Science, 18:407–419.

Kunte, K. 2008. Mimetic butterflies support Wallace’s model of sexual dimorphism. Proceedings of the Royal Society B, 275:1617–1624.

Kunte, K. 2009. Female-limited mimetic polymorphism: A review of theories and a critique of sexual selection as balancing selection. Animal Behaviour, 78:1029–1036.

Kunte, K., A. G. Kizhakke, and V. Nawge. 2021. Evolution of mimicry rings as a window into community dynamics. *Annual Review of Ecology*, Evolution, and Systematics, 52:315–341.

Kunte, K., S. Sondhi, and P. Roy. 2026. Butterflies of India, v. 4.18. Retrieved February 25, 2026. url: https://www.ifoundbutterflies.org.

Larsen, E. A., M. W. Belitz, R. P. Guralnick, and L. Ries. 2022. Consistent trait-temperature interactions drive butterfly phenology in both incidental and survey data. Scientific Reports, 12:13370.

Long, E. C., K. F. Edwards, and A. M. Shapiro. 2015. A test of fundamental questions in mimicry theory using long-term datasets. Biological Journal of the Linnean Society, 116:487–494.

Mappes, J., H. Kokko, K. Ojala, and L. Lindström. 2014. Seasonal changes in predator community switch the direction of selection for prey defences. Nature Communications, 5:5016.

Murali, K. S. and R. Sukumar. 1993. Leaf flushing phenology and herbivory in a tropical dry deciduous forest, southern India. Oecologia, 94:114–119.

Navarro-Cano, J. A., B. Karlsson, D. Posledovich, T. Toftegaard, C. Wiklund, J. Ehrlén, and K. Gotthard. 2015. Climate change, phenology, and butterfly host plant utilization. AMBIO, 44:78–88.

Pai, D., M. Rajeevan, O. . Sreejith, B. Mukhopadhyay, and N. . Satbha. 2014. Development of a new high spatial resolution (0.25° × 0.25°) long period (1901-2010) daily gridded rainfall data set over India and its comparison with existing data sets over the region. MAUSAM, 65:1–18.

Peterson, M. A. 1997. Host Plant Phenology and Butterfly Dispersal: Causes and Consequences of Uphill Movement. Ecology, 78:167.

Posledovich, D., T. Toftegaard, C. Wiklund, J. Ehrlén, and K. Gotthard. 2018. Phenological synchrony between a butterfly and its host plants: Experimental test of effects of spring temperature edited by C. Teplitsky. Journal of Animal Ecology, 87:150–161.

Prudic, K. L., B. N. Timmermann, D. R. Papaj, D. B. Ritland, and J. C. Oliver. 2019. Mimicry in viceroy butterflies is dependent on abundance of the model queen butterfly. Communications Biology, 2:68.

Repetto, M. F., M. E. Torchin, G. M. Ruiz, C. Schlöder, and A. L. Freestone. 2024. Biogeographic and seasonal differences in consumer pressure underlie strong predation in the tropics. Proceedings of the Royal Society B: Biological Sciences, 291.

Ries, L. and S. P. Mullen. 2008. A rare model limits the distribution of its more common mimic: A twist on frequency-dependent Batesian mimicry. Evolution, 62:1798–1803.

Sam, K., E. Sivault, S. Fernandez Garzon, S. Finnie, J. Kollross, M. Houska Tahadlova, J. Lenc, M. Libra, A. Ludwig, H. Maraia, A. J. Philip, L. Re Jorge, X. Xiao, and M. Volf. 2026. Forest canopy insects are safer from predators in the tropics than at higher latitudes. Nature Communications, 17:3283.

Stouffer, P. C., E. I. Johnson, and R. O. Bierregaard. 2013. Breeding seasonality in central Amazonian rainforest birds. The Auk, 130:529–540.

Su, S., M. Lim, and K. Kunte. 2015. Prey from the eyes of predators: Color discriminability of aposematic and mimetic butterflies from an avian visual perspective. Evolution, 69:2985–2994.

Sullivan, B. L., C. L. Wood, M. J. Iliff, R. E. Bonney, D. Fink, and S. Kelling. 2009. eBird: A citizen-based bird observation network in the biological sciences. Biological Conservation, 142:2282–2292.

Toftegaard, T., D. Posledovich, J. A. Navarro Cano, C. Wiklund, K. Gotthard, and J. Ehrlén. 2019. Butterfly–host plant synchrony determines patterns of host use across years and regions. Oikos, 128:493–502.

Tsurui Sato, K., Y. Sato, E. Kato, M. Katoh, R. Kimura, H. Tatsuta, and K. Tsuji. 2019. Evidence for frequency dependent selection maintaining polymorphism in the Batesian mimic Papilio polytes in multiple islands in the Ryukyus, Japan. Ecology and Evolution, 9:5991–6002.

Waldbauer, G. P. 1988. Asynchrony between Batesian mimics and their models. American Naturalist, 131 Suppl:S103--S121.

Waples, R. S. 2025. The Idiot’s Guide to Effective Population Size. Molecular Ecology, 34.

Wolda, H. 1988. Insect seasonality: Why? Annual Review of Ecology and Systematics, 19:1–18.

Wood, S. N. 2011. Fast stable restricted maximum likelihood and marginal likelihood estimation of semiparametric generalized linear models. Journal of the Royal Statistical Society Series B: Statistical Methodology, 73:3–36.

Zvereva, E. L. and M. V. Kozlov. 2021. Seasonal variations in bird selection pressure on prey colouration. Oecologia, 196:1017–1026.

